# Spin-State Modulation by Atom–Cluster Synergy Steers H2O2 Conversion toward a Catalase-like Decomposition Pathway for Anti-Inflammatory Therapy

**DOI:** 10.64898/2026.03.01.708785

**Authors:** Hanjie Zhang, Xiaoli Wang, Qi Sun, Dongze Mo, Shujie Liu, Wanling Liu, Xiaomiao Cui, Xueying An, Jiang Du, Zhenzhen Wang, Xingfa Gao, Hui Wei

**Author notes:** These authors contributed equally to this work (Hanjie Zhang, Xiaoli Wang).

## Abstract

Nanozymes have emerged as promising enzyme mimics for anti-inflammatory therapy. However, their catalytic efficiency and substrate selectivity generally remain inferior to those of natural enzymes. Single-atom nanozymes (SAzymes), with isolated metal centers resembling enzymatic active sites, represent an important advance toward rational design of nanozyme, but achieving enzyme-like selectivity remains challenging. Herein, we reported a spin-state modulation strategy to prepare Fe–N–C nanozyme with coexisting single atoms and nanoclusters (Fe_SA+NC_) via reductive-gas pyrolysis. Experimental analyses and density functional theory calculations revealed that Fe nanoclusters induced local symmetry breaking and charge redistribution around FeN_4_ sites, shifting the Fe centers toward a higher spin configuration and thereby modulating the free energy changes of the H_2_O_2_ conversion pathway. As a result, Fe_SA+NC_ showed dramatically enhanced catalase (CAT)-like activity (333.79 U mg^−1^) and suppressed peroxidase (POD)-like activity (38.49 U mg^−1^), achieving superior selectivity compared to Fe SAzymes (Fe_SA_), which showed comparable CAT- and POD-like activities (62.39 and 60.39 U mg^−1^, respectively). Moreover, Fe_SA+NC_ achieved a higher superoxide dismutase (SOD)-like activity (929.27 U mg^−1^) than Fe_SA_ (249.68 U mg^−1^), enabling efficient SOD-CAT cascade. Fe_SA+NC_ effectively scavenged excessive intracellular reactive oxygen species, suppressed M1 macrophage polarization, and enhanced the therapeutic efficacy of intra-articular stem cell injection in a rat model of rheumatoid arthritis, a representative chronic inflammatory disease. This work highlights an atom–cluster synergy strategy for steering H_2_O_2_ conversion towards antioxidant pathway, offering a general design principle for safer, more controllable and more efficient nanozyme-based anti-inflammatory therapeutics.

## Introduction

Chronic inflammatory, such as rheumatoid arthritis (RA), are often accompanied by excessive production of reactive oxygen species (ROS). ^[1,2]^ In inflammatory tissues, cells continuously generate superoxide anions (O_2_^•−^), hydroxyl radicals (^•^OH), and hydrogen peroxide (H_2_O_2_), leading to lipid peroxidation, mitochondrial injury, and irreversible tissue damage. ^[3–5]^ In physiological antioxidant systems, multiple enzymes cooperate to regulate ROS levels. Superoxide dismutase (SOD) catalyzes the disproportionation of O_2_^•−^ into O_2_ and H_2_O_2_, while catalase (CAT) subsequently decomposes H_2_O_2_ into O_2_ and H_2_O through a non-radical dismutation pathway, effectively terminating the expansion of oxidative damage. ^[4,6,7]^ Although enzymes possess superior catalytic efficiency and substrate specificity, single enzyme intervention is often insufficient for restoring redox homeostasis in complex ROS networks, which normally require synergistic action of multiple antioxidant enzymes.

Nanozymes, owing to their multifunctionality and robust physicochemical stability, provide an attractive alternative to natural enzymes. ^[8]^ By integrating multiple enzyme-like activities into a single material, nanozymes could realize physiological SOD-CAT cascade and achieve continuous ROS scavenging. ^[9]^ However, the coexistence of these activities is often poorly controlled, particularly in the way nanozymes activate H_2_O_2_ through CAT-like and peroxidase (POD)-like pathways. ^[10]^ This leads to compromised selectivity and undesired radical generation, which limits their therapeutic safety and efficacy *in vivo*.

Single-atom nanozymes (SAzymes) feature atomically dispersed metal centers with well-defined coordination environments and fully exposed active sites, enabling high catalytic efficiency comparable to natural enzymes. ^[11,12]^ Recent studies have employed SAzymes to achieve precise regulation of enzyme-like activity. ^[13–15]^ Nevertheless, symmetric metal–N_4_ construction motif in typical SAzymes may rigidify the local electronic structure and constrain the adsorption and activation of reaction intermediates, thereby limiting fine control over H_2_O_2_ conversion pathways. ^[16–18]^ Efficient SAzyme design hinges on optimizing the coordination environment of the catalytic site and elucidating structure-activity relationships. To modulate the electronic structure and catalytic activity, several approaches have been explored, including varying the metal coordination number; ^[12,19]^ introducing heteroatom ligands (P, S, O, B, Cl, etc.); ^[11,15,17,20–22]^ incorporating secondary metal centers; ^[23,24]^ and constructing hybrid systems in which single atoms coexist with nanoclusters or nanoparticles. ^[25,26]^ In such atom–cluster ensembles, electron redistribution between components establishes a charge-transfer gradient that modulates adsorbate bonding configurations and reaction pathways. ^[27,28]^ Moreover, the unoccupied d orbitals of single-atom sites can couple with the s/p/d orbitals of the nanoclusters, leading to orbital hybridization and shifts in energy levels. ^[25,29]^ These interactions adjust the Fermi level of the active sites and tune the binding strength of key intermediates, ^[29,30]^ thereby regulating reaction selectivity and activation barriers. ^[31,32]^

Herein, we reported an Fe-based atom–cluster composite nanozyme (Fe_SA+NC_) featuring coexisting Fe single atoms and ultrasmall Fe nanoclusters, prepared via reductive-gas-assisted pyrolysis. Different from conventional Fe SAzymes (Fe_SA_), Fe nanoclusters in Fe_SA+NC_ induced local symmetry breaking and charge redistribution around FeN_4_ sites. Atom–cluster synergy drove Fe centers towards a higher spin configuration and modulated the free energy changes of the H_2_O_2_ conversion pathway. Consequently, Fe_SA+NC_ exhibited a markedly CAT-like activity (333.79 U mg^−1^) with rather restrained POD-like activity (38.49 U mg^−1^), and achieved a high catalytic kinetic V_max_ value (0.39 mM s^−1^), superior to those of Fe_SA_ and most reported nanozymes. Fe_SA+NC_ achieved significantly improved reaction selectivity toward antioxidant H_2_O_2_ decomposition. By contrast, Fe_SA_ exhibited comparable CAT- and POD-like activities (62.39 and 60.39 U mg^−1^, respectively). Moreover, the SOD-like activity of Fe_SA+NC_ (929.27 U mg^−1^) exceeded that of Fe_SA_ (249.68 U mg^−1^), enabling efficient SOD-CAT cascade for ROS scavenging. As a result, Fe_SA+NC_ performed excellent antioxidant and anti-inflammatory efficacy both *in vitro* and *in vivo*. Using a rat model of RA as a proof-of-concept, Fe_SA+NC_ scavenged excessive intracellular ROS, suppressed pro-inflammatory (M1) macrophage polarization, and enhanced the therapeutic efficacy of intra-articular stem cell injection in RA rats. These findings suggest that atom–cluster synergy may serve as a promising design strategy to modulate H_2_O_2_ conversion selectivity. By steering H_2_O_2_ decomposition toward a CAT pathway while suppressing radical-generating routes, such a strategy is crucial for reducing harmful ROS production and improving the safety and controllability of nanozyme-based anti-inflammatory therapeutics.

**Scheme 1.**
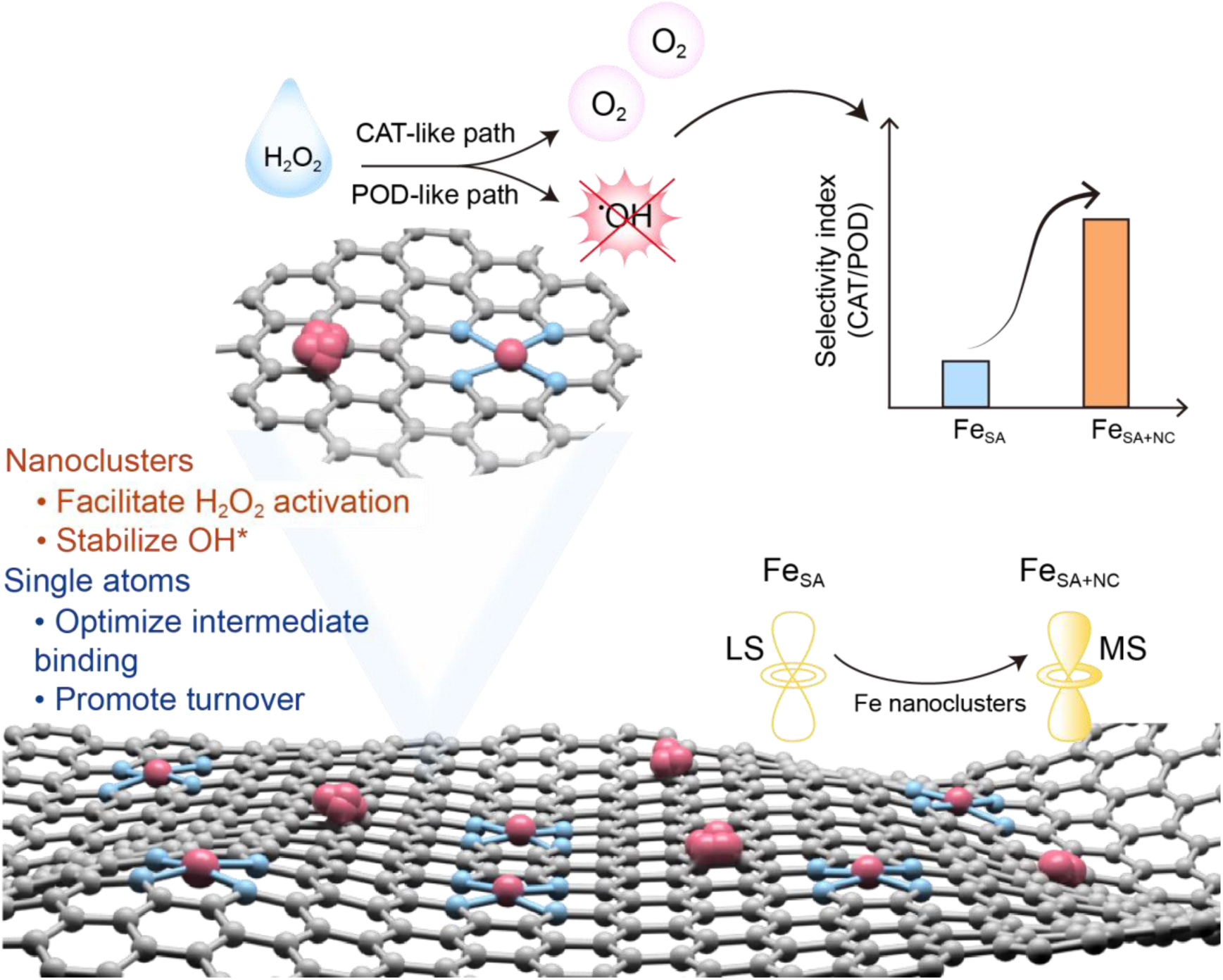
Atom–cluster synergy in Fe_SA+NC_ tunes the electronic structure to favor CAT-like H_2_O_2_ decomposition (O_2_ evolution) while suppressing POD-like ^•^OH formation, yielding a higher CAT/POD ratio than Fe_SA_.

## Results

### Synthesis and structural characterizations of Fe**–**N**–**C nanozymes

The preparation of Fe_SA+NC_ and Fe_SA_ were demonstrated schematically in Figure 1a. Fe(acac)_3_ was first encapsulated in zeolitic imidazolate framework-8 (ZIF-8) to form Fe@ZIF-8. The X-ray diffraction (XRD) pattern of Fe@ZIF-8 matched with that of pristine ZIF-8 (Figure S1a). Fe@ZIF-8 was then pyrolyzed at 900 ℃ under H_2_/Ar to obtain Fe_SA+NC_, or under Ar to obtain Fe_SA_. After pyrolysis, both samples showed similar XRD patterns with no detectable metal or metal-oxide peaks. The two broad shoulder peaks around 23° and 44° were assigned to the carbon (002) and (101) diffraction (Figure S1b). Transmission electron microscopy (TEM) imaging revealed that both Fe_SA+NC_ and Fe_SA_ maintained the regular dodecahedron of ZIF-8 after pyrolysis treatment (Figure S2). Moreover, using aberration-corrected high-angle annular dark field-scanning transmission electron microscopy (HAADF-STEM) imaging, the atomic structures of Fe_SA+NC_ and Fe_SA_ were analyzed. Fe_SA_ showed only isolated bright spots (yellow circles highlighted), indicating atomically dispersed Fe single atoms without nanoclusters (Figure 1b). In contrast, Fe_SA+NC_ contained abundant Fe single atoms together with Fe nanoclusters (blue circles highlighted, Figure 1d). Energy-dispersive spectroscopy (EDS) mappings further confirmed the presence of Fe, N, and C in both samples (Figure 1c and 1e). Despite the presence of nanoclusters, Fe_SA+NC_ showed no crystalline diffraction peaks in XRD, suggesting that the nanoclusters were very small and the carbon framework remained dominant. ^[33,34]^

**Figure 1.**
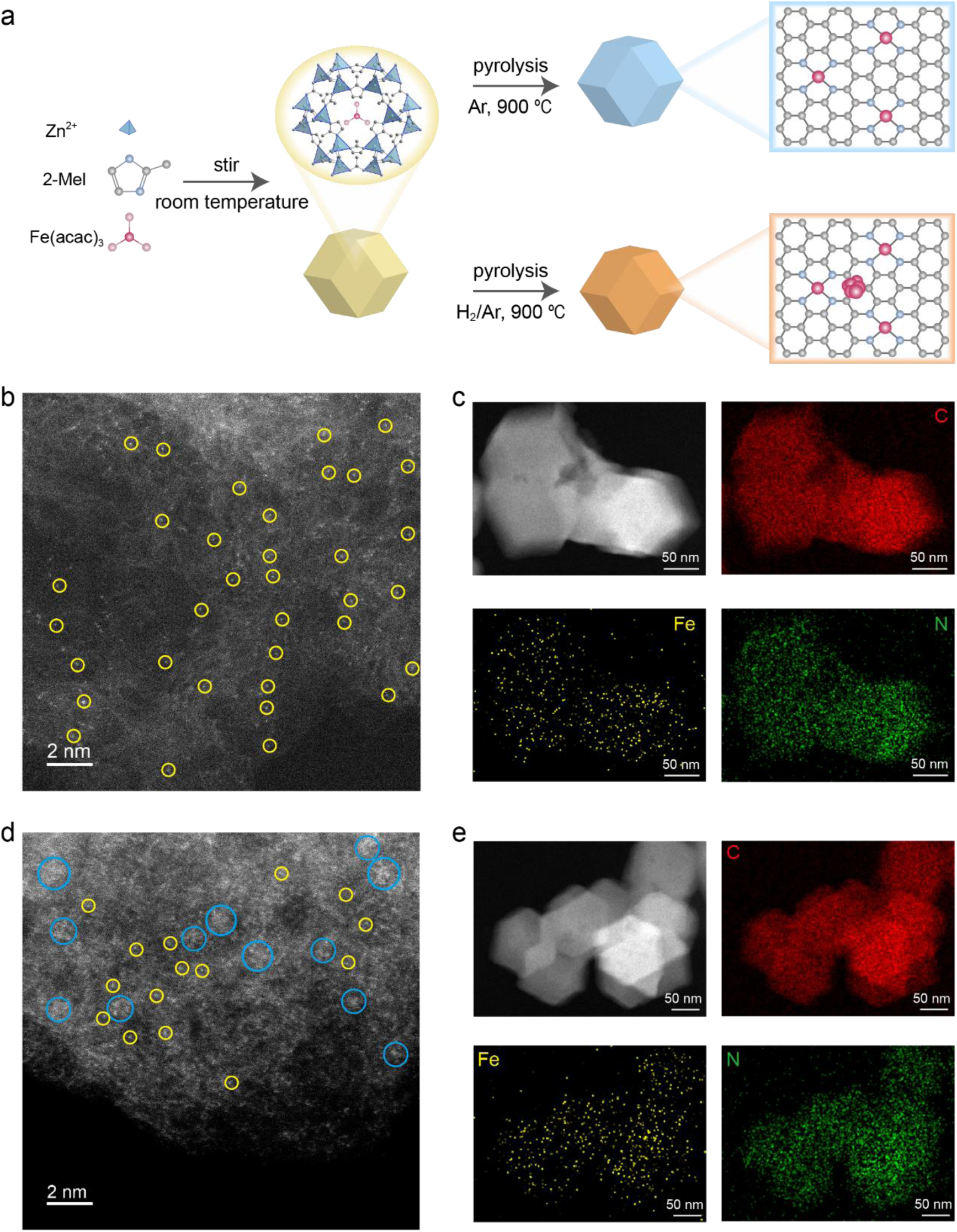
Synthesis and structural characterizations of Fe–N–C nanozymes. a) Schematic diagram of the preparation of Fe_SA_ and Fe_SA+NC_. b) HAADF-STEM image and c) the corresponding EDS mapping images of Fe_SA_. d) HAADF-STEM image and e) the corresponding EDS mapping images of Fe_SA+NC_. (Single atom, yellow circle; nanocluster, blue circle).

Brunauer–Emmett–Teller (BET) analysis displayed that Fe_SA+NC_ had a larger specific surface area and average pore volumes than those of Fe_SA_ (Figure S3 and Table S1). This contributed to the exposure of more active sites and accelerated mass transfer. Raman spectra further indicated that Fe_SA+NC_ had a lower I_D_/I_G_ ratio, suggesting a more graphitized and better-ordered carbon structure than Fe_SA_ (Figure 2a). X-ray photoelectron spectroscopic (XPS) confirmed that both samples contain C, N, and O (Figure S4a and S4b). C 1s spectra were similar for the two samples (Figure S4c and S4d), but semi-quantitative fitting showed that Fe_SA+NC_ had more C=C (∼284.6 eV) and less C–N/C–C (∼285.6 eV) (Table S2). ^[35,36]^ N 1s spectra were deconvoluted into pyridinic (∼398.5 eV), pyrrolic (∼399.5 eV), graphitic (∼401.1 eV), O-pyridine (∼402.4 eV), oxidized (∼404.4 eV), and adsorbed (∼406.1 eV) nitrogen species (Figure S4e and S4f). ^[37,38]^ As shown in Figure 2b and Table S3, Fe_SA+NC_ contained more graphitic N. Consistent with reports that the pyrolysis atmosphere can tune local C/N coordination, ^[39–41]^ H_2_/Ar pyrolysis enriched graphitic N, which could maintain electron conjugation and accelerate electron transfer during catalysis. ^[42–44]^ Fe content was quantified by using inductively coupled plasma (ICP). Three batches of samples showed similar results, confirming good reproducibility (Table S4). Besides, Fe_SA+NC_ exhibited negligible changes in hydrodynamic diameter and polydispersity index (PDI) over 72 h, indicating a narrow size distribution and the absence of appreciable aggregation or precipitation (Figure S5).

**Figure 2.**
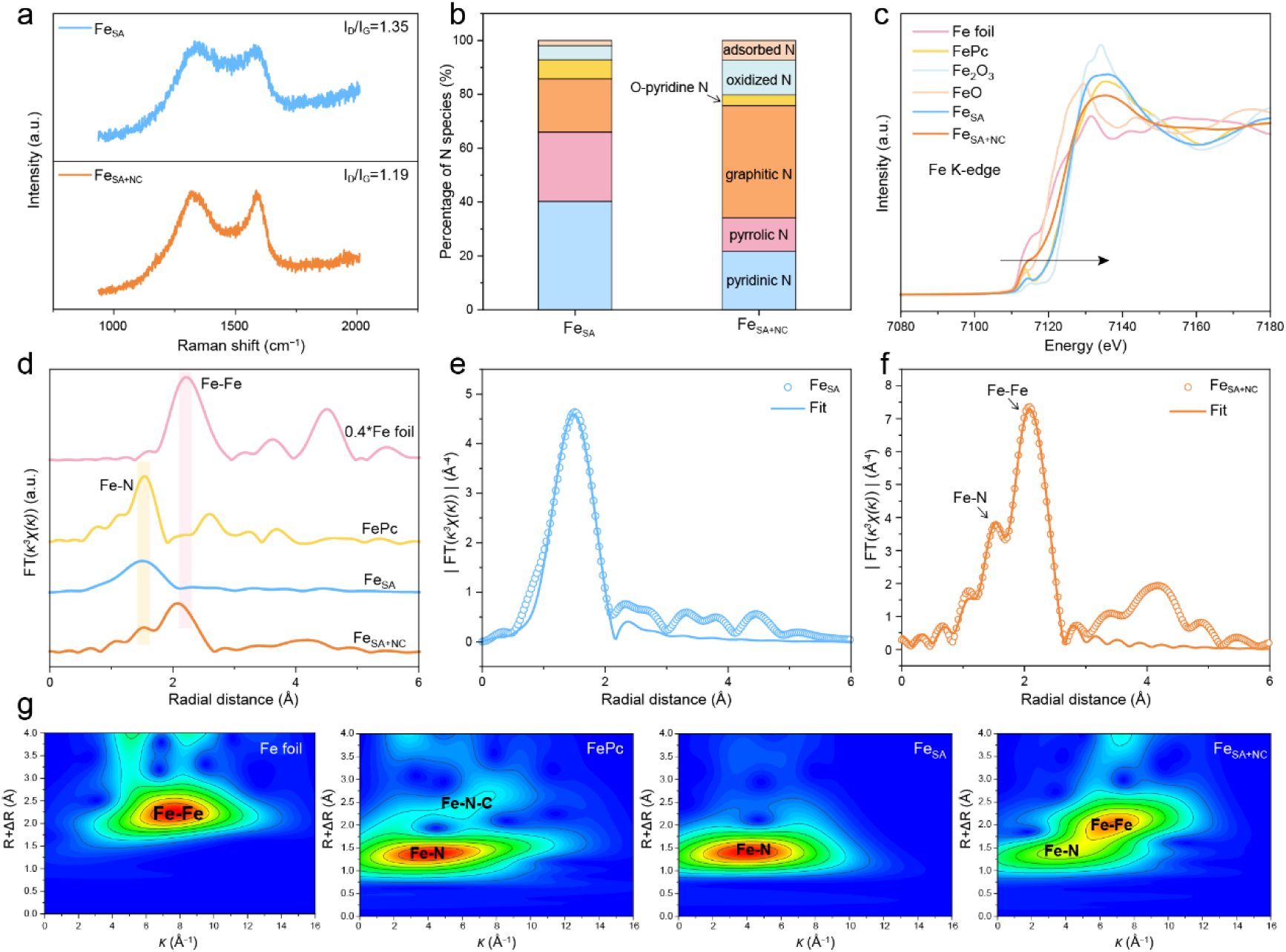
Structure analyses of Fe–N–C nanozymes. a) Raman spectra and b) N species content calculated from high-resolution N 1s XPS spectra of Fe_SA_ and Fe_SA+NC_. c) Normalized Fe K-edge XANES spectra, d) *κ^3^*-weighted FT-EXAFS spectra and g) wavelet transforms of Fe foil, FePc, Fe_SA_, and Fe_SA+NC_. Fe K-edge experimental and fitting curves of e) Fe_SA_ and f) Fe_SA+NC_ in R space.

### Atomic structure analyses of Fe**–**N**–**C nanozymes

The coordination structures and bonding configurations of Fe_SA+NC_ and Fe_SA_ were investigated using the synchrotron-based X-ray absorption near edge structure (XANES) and the extended X-ray absorption fine structure (EXAFS) spectra. As shown in Figure 2c, the absorption edges of both samples lay between Fe foil and Fe_2_O_3_, indicating partially oxidized Fe. Linear fitting analysis using Fe foil, FeO, and Fe_2_O_3_ as references, the average Fe valence was 2.92 for Fe_SA_ and 2.59 for Fe_SA+NC_ (Figure S6). Compared with Fe_SA_, Fe_SA+NC_ showed a markedly reduced white-line peak intensity at ∼7135 eV. As Fe K-edge white-line peak originated from the transition of electrons from the 1s to the unoccupied 4p states, a weaker intensity meant a few empty orbitals for electrons. ^[11]^ This may be due to the presence of Fe nanoclusters, which led to a decrease in the oxidation state and an increase in the number of d-electrons of Fe atom in Fe_SA+NC_. Besides, pre-edge peaks at ∼7114 eV were observed in Fe_SA+NC_, Fe_SA_, and iron phthalocyanine (FePc), indicating the existence of square-planar FeN_4_ configuration similar to porphyrin structure. ^[29]^ However, Fe_SA+NC_ showed an enhanced and broadened pre-edge feature, which suggested a disorder in highly symmetric square geometry around Fe. ^[25,45]^ Such reduced local symmetry strengthened 3d–4p mixing, thereby promoting the transition from 1s to 3d. ^[46]^ Furthermore, the *κ*^3^-weight Fourier transform EXAFS (FT-EXAFS) were analyzed to characterize their coordination structure (Figures 2d-2f, S7, and S8). Fe_SA_ exhibited an obvious peak at around 1.5 Å, which was corresponded to the Fe–N first coordination shell and the fitted coordination number was 4.1 (Figure 2e and Table S5). In addition to Fe–N peak, Fe_SA+NC_ exhibited another distinct peak at around 2.1 Å, indicating the presence of a Fe–Fe contribution. This assignment was supported by wavelet transform (WT) analysis (Figure 2g) and the HAADF-STEM observations (Figure 1d). Notably, the Fe–Fe peak in Fe_SA+NC_ was slightly shifted to lower R compared with Fe foil (2.2 Å), which likely reflected the different local coordination environment and the interaction between Fe nanoclusters and Fe–N sites. The fitting result of EXAFS represented the coordination number of Fe–N and Fe–Fe in Fe_SA+NC_ were 3.9 and 2.7, respectively (Figure 2f and Table S5). With Fe foil and FePc as references, WT contour plots showed that Fe_SA_ exhibited only Fe–N scattering path, while Fe_SA+NC_ displayed both Fe–Fe and Fe–N scattering paths (Figure 2g). Overall, Fe_SA_ was dominated by isolated FeN_4_ site, while Fe_SA+NC_ contained FeN_4_-like sites together with small Fe nanocluster. The evolution from single atoms to atom–cluster hybrid has the potential to overcome the linear scaling limitation and precisely tune the adsorption energies of various reaction intermediates. Therefore, the catalytic system may achieve superior performance towards different substrates.

### CAT-like activity analyses of Fe**–**N**–**C nanozymes

CAT is a key antioxidant enzyme that covert H_2_O_2_ into H_2_O and O_2_, thereby protecting organisms from oxidative stress. To evaluate the CAT-like activity of Fe_SA_ and Fe_SA+NC_, the O_2_ generation were monitored using a dissolved oxygen meter. As shown in Figure S9, Fe_SA+NC_ produced O_2_ more rapidly than Fe_SA_. Specific CAT-like activities were further calculated quantitatively by tracking the decrease of ultraviolet (UV) absorbance for H_2_O_2_ at 240 nm. The consumption rate of H_2_O_2_ increased with reaction time and catalyst concentration (Figure 3a and 3b). Notably, Fe_SA+NC_ achieved a specific activity of 333.79 ± 16.65 U mg^−1^, which was 5.35 times that of Fe_SA_ (62.39 ± 1.21 U mg^−1^) as shown in Figure 3c. Kinetic parameters were then determined by changing the concentration of H_2_O_2_ (2.5–40 mM) and plotting the corresponding initial velocities against the concentrations in a reciprocal method (Figure 3d and 3e). The maximal velocities (*V*_max_) and Michaelis constant (*K*_m_) could be obtained based on the linear fitting. As shown in Figure S10a, though Fe_SA+NC_ (42.70 mM) exhibited a 2.26-fold higher *K*_m_ value than Fe_SA_ (18.92 mM), Fe_SA+NC_ seemed to be closer to natural enzyme (45.91 or 52.14 mM). ^[47,48]^ The *V*_max_ value of Fe_SA+NC_ (0.39 mM s^−1^) was 7.80-fold higher than that of Fe_SA_ (0.05 mM s^−1^), achieving a substantial activity improvement (Figure 3f). Considering that H_2_O_2_ levels in pathological environment are at micromolar level, the reaction efficiency of nanozymes at extremely low substrate concentration is of practical significance. ^[49,50]^ As shown in Figure S10, although the affinity of Fe_SA+NC_ was lower than that of Fe_SA_, the catalytic rate constant (*k*_cat_) of Fe_SA+NC_ (237.00 s^−1^) exhibited about 2.49-fold higher than that of Fe_SA_ (95.12 s^−1^). We also monitored the intrinsic CAT-like activity of nitrogen doped carbon supports (N–C_SA_ and N–C_SA+NC_). Both supports showed negligible activity (Figure S11), indicating that the active centers were only Fe atoms.

**Figure 3.**
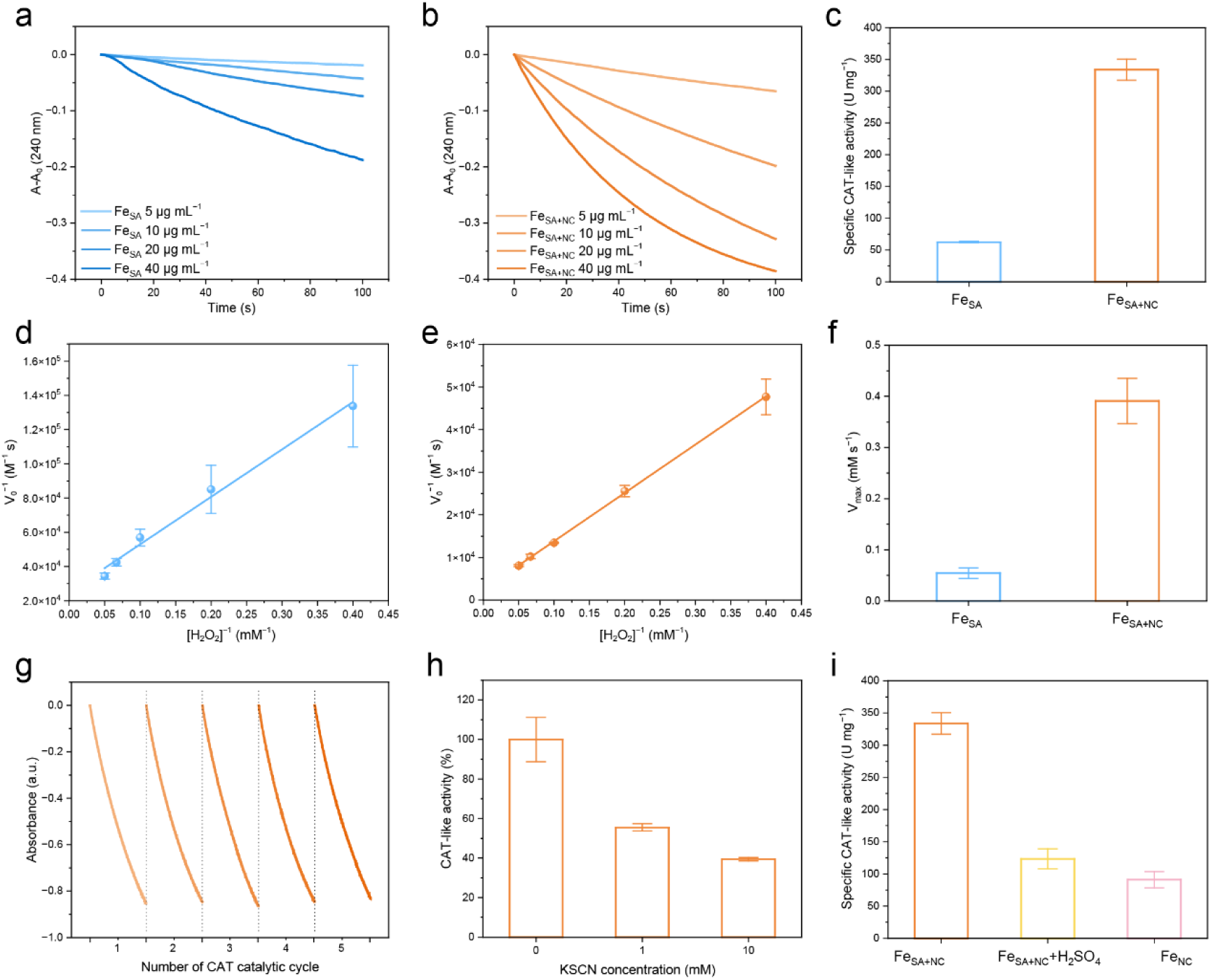
CAT-like activity and kinetics analyses of Fe–N–C nanozymes. a) Time-dependent UV absorbance at 240 nm for monitoring the CAT-mimicking catalytic activities of a) Fe_SA_ and b) Fe_SA+NC_. c) Comparison of specific CAT-like activity of Fe_SA_ and Fe_SA+NC_. Double-reciprocal plots of initial velocities versus reciprocal the concentrations of H_2_O_2_ of d) Fe_SA_ and e) Fe_SA+NC_. f) Comparison of V_max_ values of Fe_SA_ and Fe_SA+NC_. g) Repetitive catalase-mimicking catalytic activity of Fe_SA+NC_. h) Changes in CAT-like activities of Fe_SA+NC_ after incubation with different concentration of KSCN. i) Specific CAT-like activity of Fe_SA+NC_, H_2_SO_4_ treated Fe_SA+NC_ and Fe_NC_. Data in c–i) are expressed as mean ± SD, n = 3.

To evaluate robustness, we measured their CAT-like activities under various pH conditions (Figure S12). The results showed that Fe_SA+NC_ maintained appreciable CAT-like activity under both acidic and alkaline environments. Surprisingly, all the specific CAT-like activities of Fe_SA+NC_ were above 200 U mg^−1^, consistently outperforming Fe_SA_. Additionally, Fe_SA+NC_ exhibited sustained catalytic performance, nearly no decrease in activity after 5 catalytic cycles (Figures 4g and S13).

**Figure 4.**
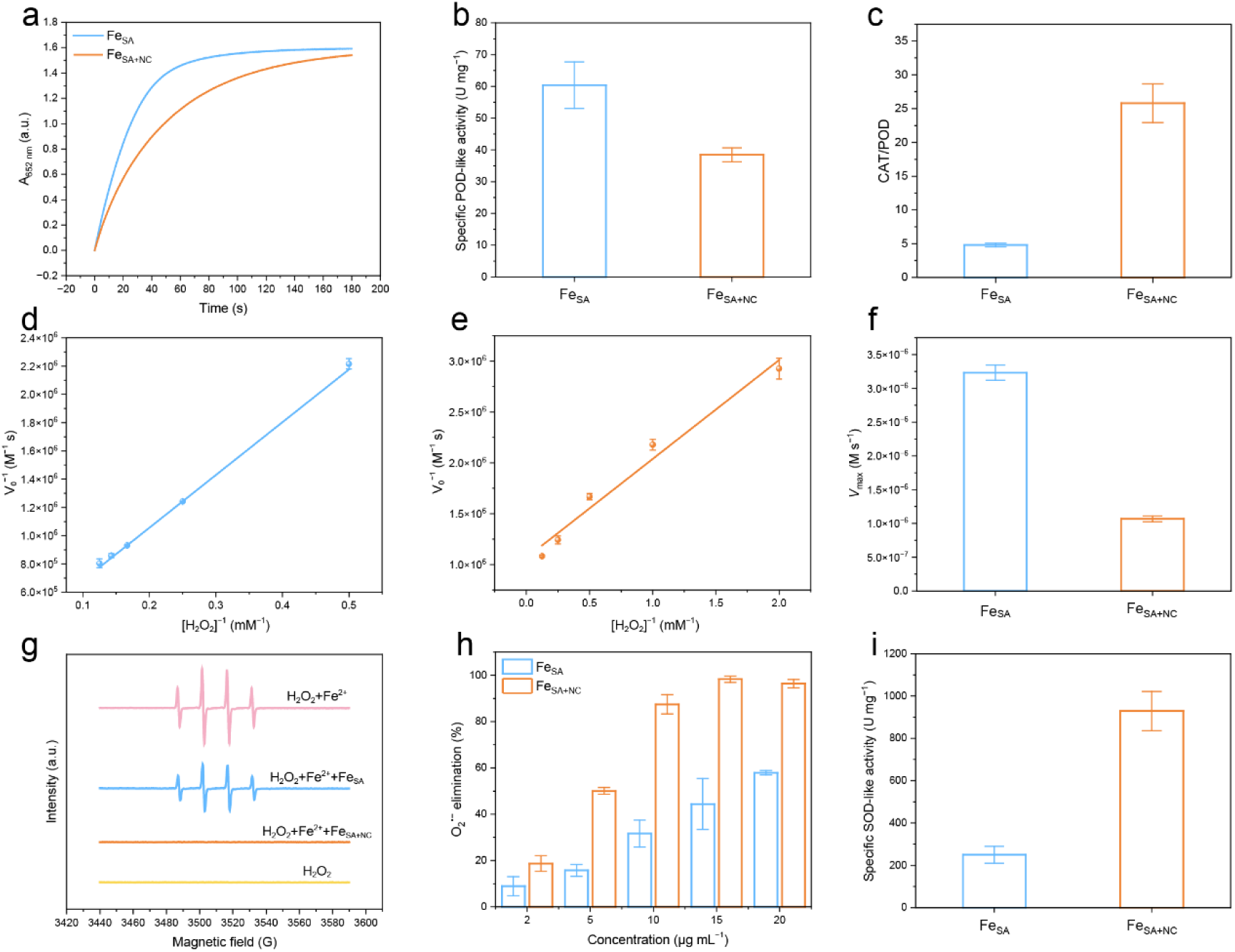
Other enzyme-like activities analyses of Fe–N–C nanozymes. a) The velocities of POD-like activity of Fe_SA_ and Fe_SA+NC_ monitored by the colorimetric reaction of TMB. b) Comparison of specific POD-like activity of Fe_SA_ and Fe_SA+NC._ c) Comparison of CAT-to-POD specific activity ratios (selectivity index) for Fe_SA_ and Fe_SA+NC_. Double-reciprocal plots of initial velocities versus reciprocal the concentrations of H_2_O_2_ of d) Fe_SA_ and e) Fe_SA+NC_. f) Comparison of V_max_ values of Fe_SA_ and Fe_SA+NC_. g) ^•^OH-scavenging ability of Fe_SA_ and Fe_SA+NC_ captured by DMPO. h) O_2_^•−^ elimination ability of Fe_SA_ and Fe_SA+NC_ measured by WST-1 assay kit. i) Comparison of SOD-like specific activities of Fe_SA_ and Fe_SA+NC_. Data in b–f and h, i) are expressed as mean ± SD, n = 3.

To further clarify the catalytic active sites, KSCN was employed as a poisoning agent to selectively block the FeN_x_ coordination sites in Fe_SA+NC_. The CAT-like activity decreased to 55% at 1 mM KSCN and 39% at 10 mM, indicating FeN_x_ were indispensable for H_2_O_2_ decomposition (Figure 3h). Then, Fe_SA+NC_ was subjected to sulfuric acid leaching to remove Fe clusters, yielding Fe SAzyme in which the atomically dispersed FeN_x_ sites were largely preserved. Fe nanocluster (Fe_NC_) was also prepared (Figure S14). As shown in Figure 3i, the H_2_SO_4_ treated Fe_SA+NC_ and Fe_NC_ both exhibited a marked decrease in CAT-like activity. These results collectively confirmed that the atomically FeN_x_ sites were the main catalytic sites responsible for the CAT-like activity of Fe_SA+NC_, whereas nanoclusters alone were insufficient but markedly promoted overall activity when coexisting with FeN_x_, supporting an atom–cluster synergy.

### POD- and OXD-like activity analyses of Fe**–**N**–**C nanozymes

Nevertheless, many nanozymes also show intrinsic POD-like activity, which may compete with and even counteract their CAT-like antioxidant function, potentially resulting in suboptimal or even detrimental effects for anti-inflammatory therapy. ^[51,52]^ To characterize the specificity of Fe_SA+NC_, we next investigated its POD-like activity, in which H_2_O_2_ acted as an oxidizing co-substrate to drive the oxidation of typical chromogenic substrates. Typically, we employed 3,3′,5,5′-tetramethylbenzidine (TMB) as a catalytic substrate. Oxidized TMB (oxTMB) showed a characteristic peak at 652 nm. According to the change of absorbance at 652 nm, the specific POD-like activities were 60.39 ± 7.31 U mg^−1^ for Fe_SA_ and 38.49 ± 2.21 U mg^−1^ for Fe_SA+NC_, indicating a 36% decrease upon introducing nanoclusters (Figure 4a and 4b). Changing the concentration of H_2_O_2_, the corresponding initial velocities were further plotted against the concentrations in a reciprocal method (Figures 4d, 4e, and S15). The *V*_max_, *K*_m_, and *k*_cat_ could be obtained based on the linear fitting (Table S6). The *V*_max_ value of Fe_SA_ (3.23 μM s^−1^) was 3.02 times that of Fe_SA+NC_ (1.07 μM s^−1^), and *k*_cat_ of Fe_SA_ (11.27 s^−1^) was 8.47 times that of Fe_SA+NC_ (1.29 s^−1^), indicating that Fe_SA_ was a much more efficient POD mimic than Fe_SA+NC_. However, high POD-like activity alone did not reflect how these nanozymes partition H_2_O_2_ between POD- and CAT-like pathways. Therefore, we defined a selectivity index as the ratio of specific CAT- to POD-like activities (Figure 4c). Fe_SA+NC_ exhibited 5.36-fold higher catalytic selectivity than that of Fe_SA_, indicating that Fe_SA+NC_ preferentially decomposed H_2_O_2_ to produce O_2_ rather than POD-like pathway. A similar trend was also observed for the ratios of *V*_max_ and *k*_cat_ values (Figure S15). To directly probe ROS generated during H_2_O_2_ conversion, we performed electron paramagnetic resonance (EPR) spectroscopy using 5,5-dimethyl-1-pyrroline N-oxide (DMPO) as a ^•^OH trapping reagent. Fe_SA_ produced a much stronger characteristic 1:2:2:1 signal than Fe_SA+NC_, indicating that Fe_SA_ favored Fenton-like pathways to generate ^•^OH, whereas Fe_SA+NC_ suppresses ^•^OH formation by redirecting H_2_O_2_ toward CAT-like decomposition (Figure S16).

As for changing TMB concentration, Fe_SA_ and Fe_SA+NC_ exhibited similar kinetic parameters, suggesting that the difference in POD-like activity mainly derived from H_2_O_2_ activation rather than TMB oxidation (Figure S17 and Table S7). To elucidate the overall redox behavior, oxidase (OXD)-like activity was subsequently examined. Fe_SA_ and Fe_SA+NC_ displayed comparable OXD-like activity, with nearly identical O_2_-driven TMB oxidation behavior (Figure S18 and Table S7). Taken together with their divergent CAT- and POD-like activities, the distinct redox behavior between Fe_SA_ and Fe_SA+NC_ mainly arose from their different partitioning of H_2_O_2_ into CAT- versus POD-like pathways rather than from differences in OXD-like activity.

### SOD-like activity and ^•^OH scavenging activity analyses of Fe**–**N**–**C nanozymes

We next examined the SOD-like activity of Fe_SA_ and Fe_SA+NC_ toward O_2_^•−^ and their capability to scavenge ^•^OH, two key steps in breaking the ROS amplification cascade in inflamed tissues. ^•^OH scavenging was investigated via EPR spectroscopy. There was no obvious signal of Fe_SA+NC_ treated group, which meant ^•^OH had been cleared away completely, while the ^•^OH of Fe_SA_ treated group was just partially cleared (Figure 4g). Then, the SOD-like activity was quantified using a commercial WST-1 assay kit. The elimination capacity of O_2_^•−^ was concentration-dependent (Figure 4h). Surprisingly, the specific SOD-like activity of Fe_SA+NC_ was 929.27 ± 92.62 U mg^−1^, which was 3.73 times that of Fe_SA_ (249.67 ± 46.06 U mg^−1^). Moreover, the elimination rate of Fe_SA+NC_ was faster than that of Fe_SA_ under the same concentration (Figure S19).

In the above sections, specific activities were normalized to the total catalyst mass (support and metal) to assess the practical activity per dosing amount. To compare the intrinsic catalytic efficiency among SAzymes with different metal loadings, the CAT- and SOD-like activities were further renormalized to the mass of metal in each sample and expressed as U mg^−1^ metal (Figure S20 and Table S8). ^[53]^ Most nanozymes fell into either a SOD-dominant or CAT-dominant region, whereas Fe_SA+NC_ lay near the upper-right diagonal, where both CAT- and SOD-like activities were high (7.26 × 10^4^ and 20.13 × 10^4^ U mg^-1^ Fe, respectively). This position highlighted the dual superiority of Fe_SA+NC_ in dismasting O_2_^•−^ and decomposing H_2_O_2_, which may be favorable for comprehensive ROS suppression for anti-inflammatory therapy.

### Electronic properties and DFT studies of Fe–N–C nanozymes

To further clarify the origin of the superior catalytic performance of Fe_SA+NC_, the electronic structure was investigated experimentally and theoretically. As shown in Figure 5a, the work function (Φ) of nanozymes were obtained by using ultraviolet photoelectron spectroscopy (UPS). The cutoff energies (E_cutoff_) of Fe_SA_ and Fe_SA+NC_ were 16.50 and 16.78 eV, giving work functions (Φ=21.22 eV-E_cutoff_) of 4.72 and 4.44 eV, respectively. The lower work function of Fe_SA+NC_ made electron donation to oxygen-containing intermediates easier, thereby facilitating faster interfacial electron transfer. The shape and position of EPR spectra can reveal the quantity and occupancy of unpaired electrons. As shown in Figure S21, Fe_SA+NC_ shown a stronger high-spin Fe(Ⅲ) species signal at g = 4.25 than Fe_SA_, and a sharp signal at g = 2.04, indicating the presence of additional paramagnetic species (e.g., Fe centers in a different ligand field and/or defect-related radicals). ^[54,55]^ To assess how Fe clusters affect the spin state in Fe_SA+NC_, the zero-field cooling temperature-dependent (ZFC-T) magnetic susceptibility (*χ*_m_) measurement was performed (Figure 5b and 5c). The effective magnetic moments (μ_eff_) of Fe_SA_ and Fe_SA+NC_ were 1.72 μ_B_ and 3.14 μ_B_, and the number of unpaired electrons (n) were calculated to be 0.99 and 2.29, respectively 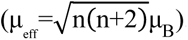. Based on the Fe valence states from XAFS (Figure S6), the electronic configuration of Fe_SA_ was d_xy_^2^d ^2^d ^1.08^ (LS, t_2g_^5.08^ e_g_ ^0^) and that of Fe_SA+NC_ was d ^2^d ^1.46^d ^1^d_z2_^1^ (MS, t ^4.46^ e_g_^1^). These results suggested that Fe nanoclusters modulated the orbital occupation and spin state of the FeN_x_.

**Figure 5.**
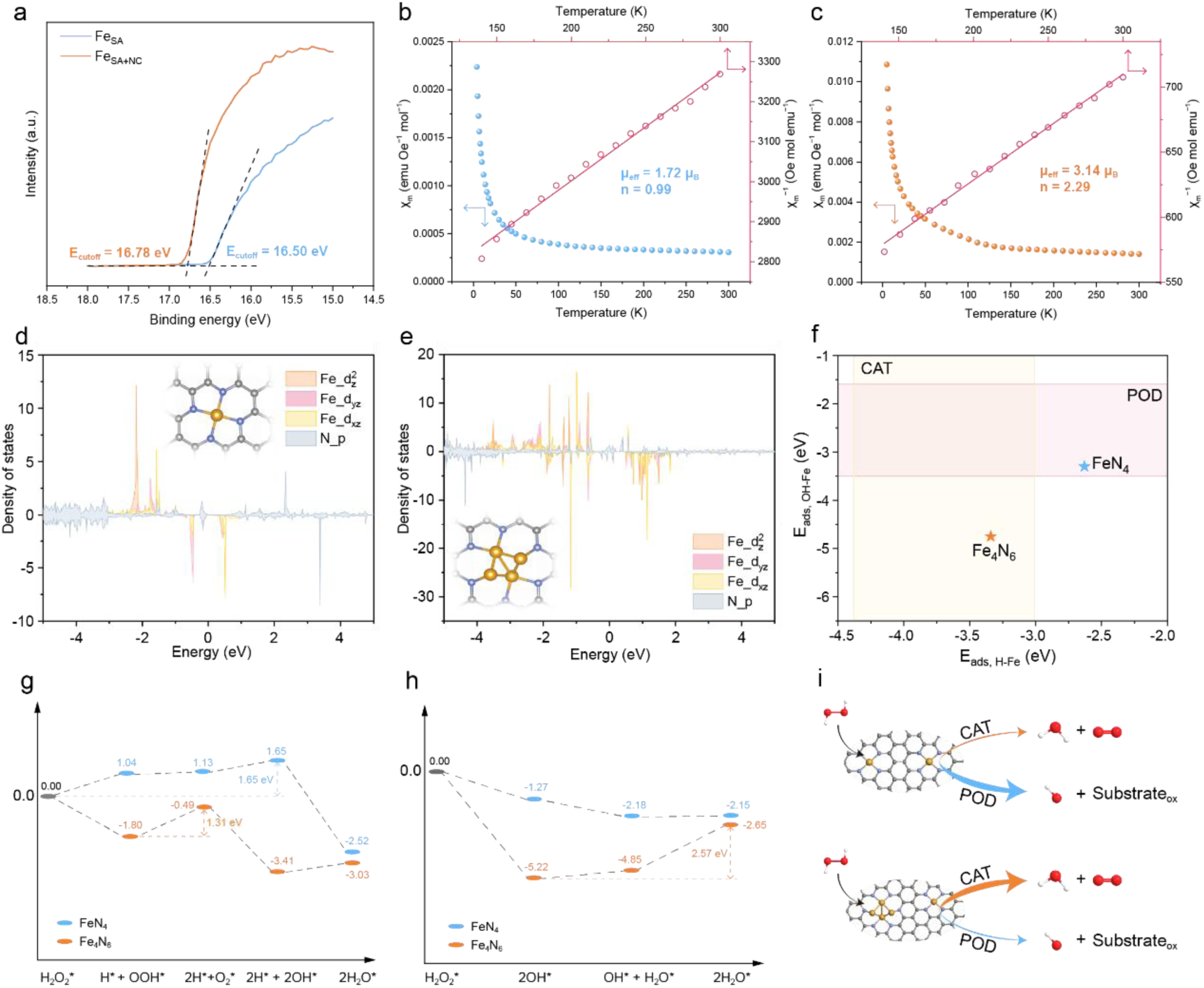
Electronic properties and theoretical analyses of Fe–N–C nanozymes. a) UPS spectra of Fe_SA_ and Fe_SA+NC_. χ_m_–T and χ ^−1^–T plots of b) Fe_SA_ and c) Fe_SA+NC_. Projected DOS diagrams of d) Fe_SA_ and e) Fe_SA+NC_. f) The hydroxyl adsorption energy (E_ads, OH_) and hydrogen adsorption energy (E_ads, H_) of FeN_4_ and Fe_4_N_6_. Free energy diagram of g) CAT-like pathway and h) POD-like pathway of FeN_4_ and Fe_4_N_6_. i) Schematic diagram of catalytic selectivity.

To elucidate the structure-property relationship, density functional theory (DFT) calculations were employed to explain how the atomic configuration influenced enzyme-like catalytic activity. The single-atom Fe and cluster Fe catalytic sites were simplified to FeN_4_ and Fe_4_N_6_, respectively. The FeN_4_ periodic model was constructed by replacing six carbon atoms in graphene with a tetrahedral FeN_4_ center, comprising one Fe atom coordinated by four N atoms. For the Fe₄N_6_ periodic model, a four-iron-atom cluster replaced a benzene ring on the graphene surface, followed by the substitution of six surrounding carbon atoms with six nitrogen atoms, forming coordination structure around the Fe cluster (Figure S22). First, projected density of states (PDOS) was calculated to assess orbital contributions and spin polarization (Figure 5d and 5e). For FeN_4_, the Fe 3d states were relatively localized and mainly distributed well below the Fermi level (E_F_), with limited density near, consistent with a less spin-polarized electronic structure. In contrast, Fe_4_N_6_ exhibited stronger spin splitting and broader Fe 3d states closer to E_F_, with evident overlap with N–p states. This indicated strengthened Fe–Fe/Fe–N electronic interactions and more spin-polarized, electronically active Fe sites. These were consistent with the stronger EPR signals and more unpaired electrons observed experimentally.

According to the descriptor framework developed by Gao and co-workers, POD-like activity is favored when hydroxyl adsorption energy (E_ads, OH_) falls into −3.5 eV to −1.6 eV, ^[56]^ while for CAT-like activity, the optimal window is either hydrogen adsorption energy (E_ads, H_) lies in −2.83 to −1.47 eV (HOMO-type) or −4.38 to −3.01 eV (LUMO-type). ^[57]^ As shown in Figure 5f, FeN_4_ had moderate OH* adsorption (−3.30 eV), which fell into POD window. In contrast, Fe_4_N_6_ over stabilized OH* (−4.75 eV), suppressing POD-like activity. Meanwhile, Fe_4_N_6_ exhibited favorable H* adsorption (−3.34 eV), located within LUMO-type CAT window.

To understand the reaction pathway, we further compared the reaction free-energy changes (Figures 5g, 5h, S23, S24 and S25). For the CAT-like route, the FeN_4_ showed an uphill profile, with the highest-energy intermediate (2H* + 2OH*) reaching 1.65 eV relative to H_2_O_2_*, indicating that H_2_O_2_ activation and O–O bond cleavage were energetically demanding on FeN_4_ sites. In contrast, the initial step (H_2_O_2_*→H* + OOH*) was exergonic on Fe_4_N_6_, suggesting easier H_2_O_2_ activation at the cluster sites. The largest uphill step along this pathway was reduced to 1.31 eV, rendering the overall H_2_O_2_ decomposition much more favorable on Fe_4_N_6_ than on FeN_4_. However, the two sites displayed opposite performance along the POD-like route. FeN_4_ proceeded downhill overall, indicating that H_2_O_2_ conversion was thermodynamically allowed through POD-like pathway on single-atom sites. In contrast, Fe_4_N_6_ showed a pronounced endergonic step toward H_2_O formation (2.57 eV), indicating that the nanocluster sites were unfavorable for converting H_2_O_2_ through POD pathway.

Together, these results revealed that Fe nanoclusters could promote CAT-like activity while suppress POD-like activity of Fe_SA+NC_ (Figure 5i). Considering the experimentally confirmed coexistence of atomically dispersed FeN_4_ sites and Fe nanoclusters in Fe_SA+NC_, we propose that the single-atom sites provide abundant, structurally well-defined redox centers, whereas the nanoclusters serve as highly active H_2_O_2_ decomposition hotspots, together steering H_2_O_2_ decomposition into a safer and more controllable pathway. Although single-atom and cluster sites were treated separately in the present calculations, they coexisted in the Fe_SA+NC_. Future models that explicitly include atom–cluster interface are expected to quantify their synergistic coupling more directly.

### Antioxidant and anti-inflammatory *in vitro*

Considering the high antioxidant activity of Fe_SA+NC_, we evaluated its potential as an anti-inflammatory nanotherapeutic medicine *in vitro*. RAW264.7 macrophages and bone marrow mesenchymal stem cells (BMSCs) were incubated with Fe_SA_ or Fe_SA+NC_ for 24 h. Fe_SA+NC_ exhibited negligible cytotoxicity, whereas Fe_SA_ caused evident cytotoxicity even at 5 μg mL^−1^ (Figure S26). Therefore, Fe_SA_ was excluded from subsequent biological experiments. We next evaluated the intracellular ROS scavenging capacity of Fe_SA+NC_. An oxidative stress model was established by treating RAW264.7 macrophages with H_2_O_2_ for 0.5 h, and ROS levels were monitored using DCFH-DA probe. Fluorescence microscopy images revealed that Fe_SA+NC_ treatment markedly reduced the green fluorescence intensity of DCFH-DA (Figure 6a), and restored the intracellular ROS level to that of the control group (Figure 6e).

**Figure 6.**
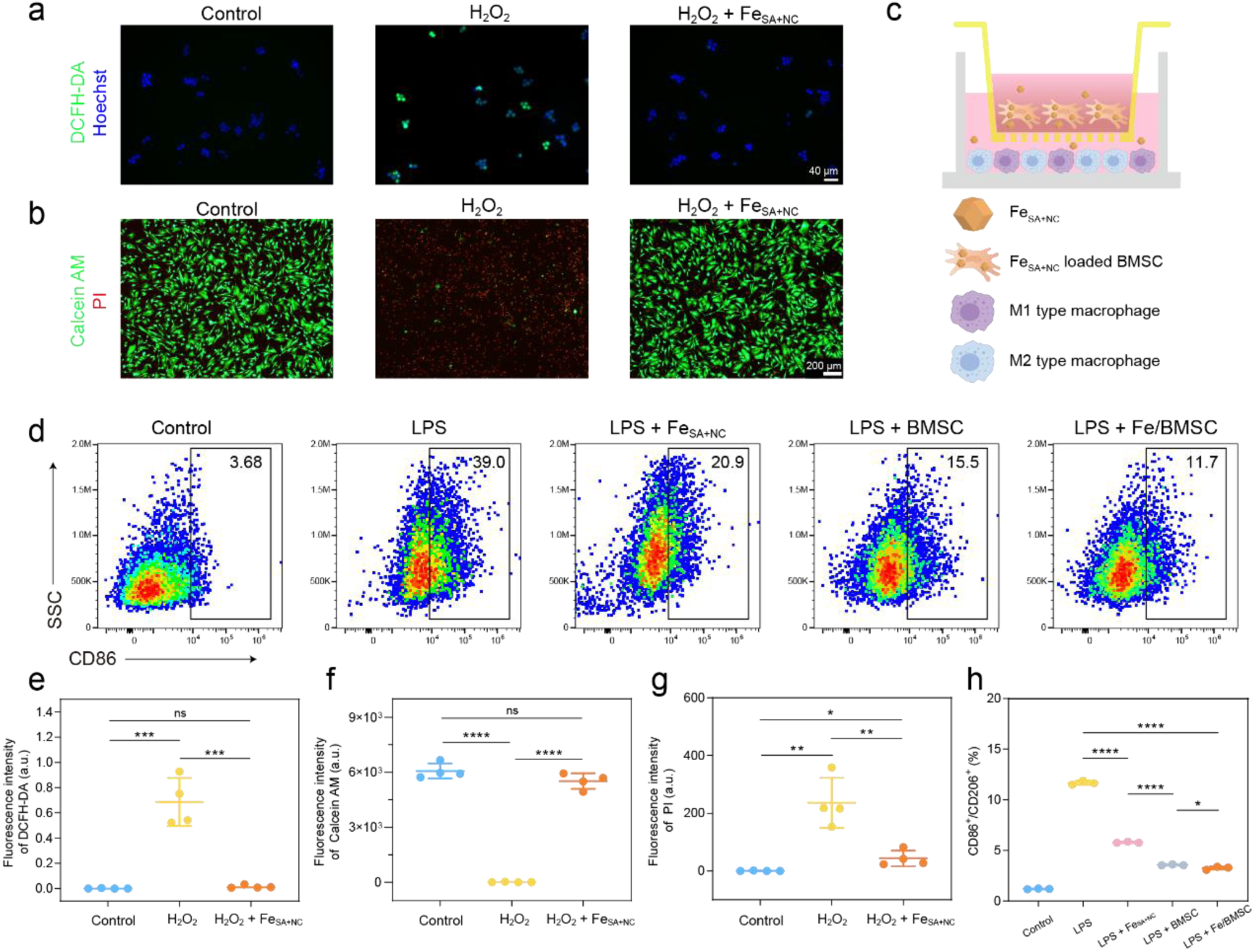
Antioxidant and anti-inflammatory ability of Fe_SA+NC_ *in vitro*. a) The ROS scavenging ability in macrophages, and b) the ability to protect BMSCs from ROS damage of Fe_SA+NC_ evaluated by fluorescence microscope. c) Schematic diagram of co-culture of BMSCs and macrophages using a Transwell system. The expression of d) CD86 in macrophages in the co-culture system evaluated by flow cytometry. Quantitative fluorescence statistics of e) ROS in macrophages, f) lived and g) dead BMSCs. h) Comparison of macrophage polarization trends in the co-culture system. Data in e–g) was presented as mean ± SD (n = 4). Data in f) was presented as mean ± SD (n = 3). Statistical differences were analyzed by one-way ANOVA and Student’s t-test: * p<0.05, ** p<0.01, *** p<0.001, and **** p<0.0001.

Prolonged exposure of cells to inflammatory microenvironments with excessive ROS compromises their viability and functional activity, a critical determinant of stem cell-based therapy. Increasing evidence indicated that stem cells engineered with antioxidant nanozymes achieve superior therapeutic outcomes. ^[58,59]^ A ROS-rich microenvironment was established by treating BMSCs with H_2_O_2_ for 24 h, and cell viability was assessed by Calcein AM/Propidium iodide (PI) staining. With Fe_SA+NC_ treatment, PI-positive cells decreased while Calcein AM-positive cells increased compared with H_2_O_2_ group (Figure 6b, 6f and 6g). These results suggested that Fe_SA+NC_ may enhance the therapeutic efficacy of administered stem cells.

Given the central roles of pro-inflammatory (M1) and anti-inflammatory (M2) macrophages in inflamed tissues, we next examined whether Fe_SA+NC_ and BMSCs formulation (Fe/BMSC) could modulate macrophage polarization (Figure 6c). Flow cytometry results showed that lipopolysaccharide (LPS) stimulation increased CD86^+^ (M1) and decreased CD206^+^ (M2) macrophages (Figures 6d and S27). In contrast, treatment with Fe_SA+NC_, BMSC or Fe/BMSC reduced the percentage of CD86^+^ macrophages. Notably, Fe/BMSC showed the strongest suppression of M1 polarization and promotion of M2 polarization (Figure 6h). This enhancement was likely owing to Fe_SA+NC_ mediated ROS scavenging and protection of BMSCs.

Collectively, *in situ* ROS scavenging by Fe_SA+NC_ may alleviate early oxidative stress during BMSC delivery, thereby supporting BMSC survival and function.

### Antioxidant and anti-inflammatory *in vivo*

RA is a progressive autoimmune disease, characterized by persistent synovitis, cartilage degeneration, and joint swelling. ^[1,2]^ Stem cell-based therapy has shown therapeutic potential for RA in numerous studies. ^[60–62]^ Meanwhile, scavenging excessive ROS and suppressing M1 macrophage polarization are recognized as effective strategies for anti-inflammation. ^[63,64]^ Based on our *in vitro* findings, we next investigated whether co-delivery of Fe/BMSC could ameliorate disease in a typical adjuvant-induced arthritis (AIA) model. Fifteen days after immunization, AIA rats were randomly divided into four groups (4 rats per group) and received intra-articular injections of PBS or PBS containing Fe_SA+NC_, BMSC or Fe/BMSC every three days, respectively (Figure 7a and 7b). Healthy rats without AIA and treatment were served as a control group. Over a 21-day treatment and observation period, the Fe/BMSC group exhibited a significant alleviation of paw swelling (Figure 7c and 7d) and the lowest arthritis scores (Figure S28).

**Figure 7.**
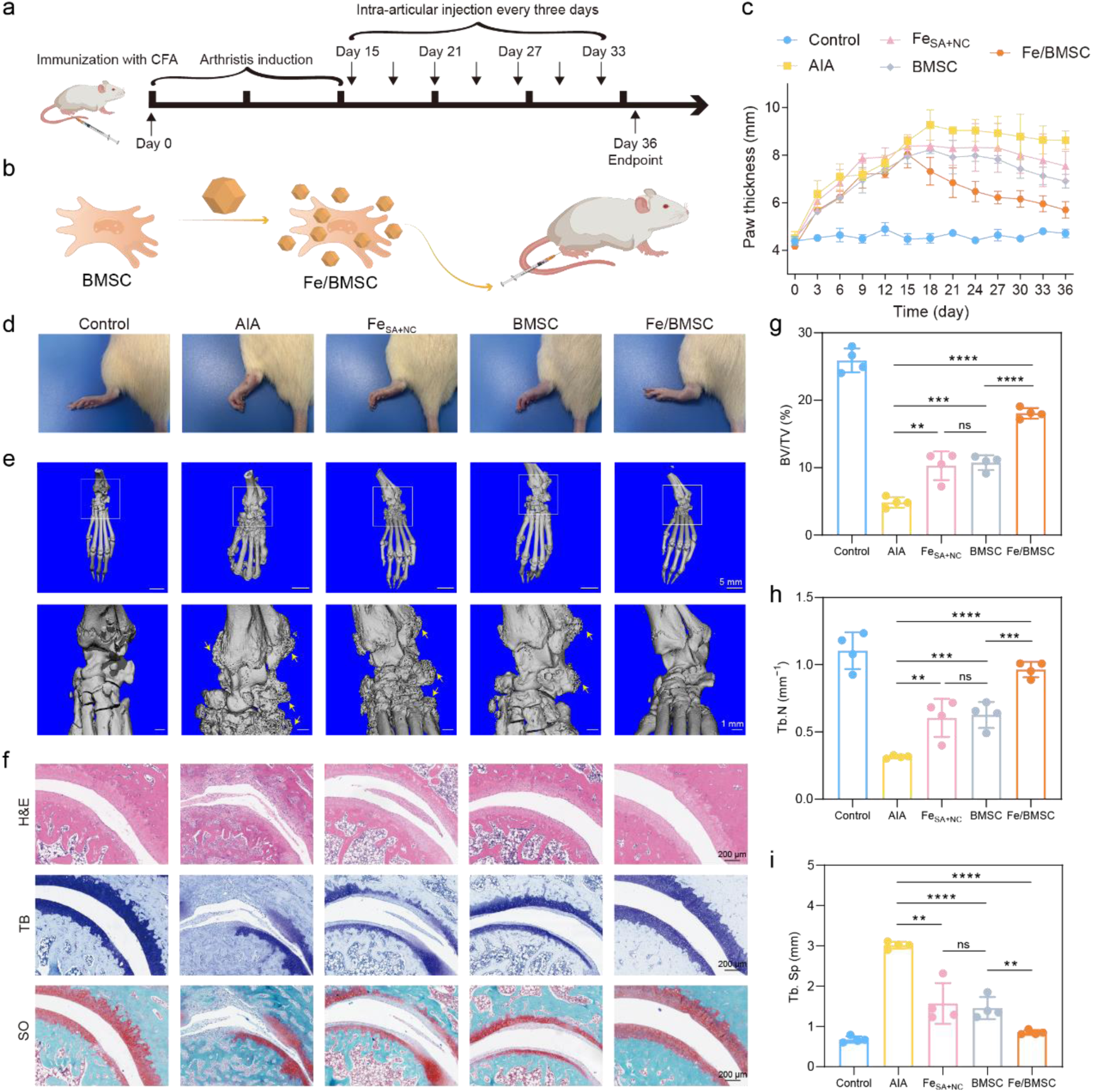
Therapeutic efficiency of Fe_SA+NC_ and BMSC in rheumatoid arthritis rat models. a) The overall timeline for rheumatoid arthritis modeling and treatment. b) Schematic diagram of co-culture of Fe_SA+NC_ and BMSC, c) The paw thickness in different groups during the treatment process. d) Representative photos and e) micro-CT images of joints of rats in different groups on day 36 and 3d reconstruction, respectively. f) Hematoxylin & eosin, toluidine blue and safranin-fixed green staining of joints after different treatments. Quantitative micro-CT analysis of g) BV/TV, h) Tb.N and i) Tb.Sp for joints of rats in different groups. Data in c, g, h and i) was presented as mean ± SD (n = 4). Statistical differences were analyzed by one-way ANOVA and Student’s t-test: * p<0.05, ** p<0.01, *** p<0.001, and **** p<0.0001.

Joint structural changes were further evaluated by micro-computed tomography (micro-CT). Compared with healthy controls, AIA rats exhibited distinct bony erosion and metatarsal swelling (Figure 7e). Treatment with Fe_SA+NC_ and BMSC alone did not afford significant improvements. In contrast, bony erosion and metatarsal swelling were largely alleviated after Fe/BMSC treatment. Micro-CT derived histomorphometric indices, including bone volume/total volume (BV/TV), trabecular number (Tb.N), and trabecular separation (Tb.Sp), indicated substantial restoration of bone microarchitecture in the Fe/BMSC group (Figure 7g–i).

Moreover, histological analysis, including hematoxylin and eosin (H&E), toluidine blue (TB), and Safranin O/Fast Green (SO) staining, further corroborated the therapeutic efficacy of Fe/BMSC (Figure 7f). Inflammatory infiltration, cartilage destruction, and synovial hyperplasia were obvious in AIA, Fe_SA+NC_, and BMSC groups. In contrast, the Fe/BMSC treated group showed smooth articular surfaces, intact cartilage structure, and minimal synovial hyperplasia. To determine macrophage polarization and quantify inflammatory cytokine levels *in vivo*, we performed immunofluorescence staining (Figure S29). In the Fe/BMSC group, the proportion of CD86⁺ macrophages was comparable to that in healthy controls, and the levels of the pro-inflammatory cytokines TNF-α and IL-1β were markedly reduced compared with the AIA group.

To assess the biosafety of Fe_SA+NC_, H&E-stained histological sections of the heart, liver, spleen, lung, and kidney are examined. No appreciable differences were observed among the four modeling groups and the health control group (Figure S30).

## Conclusion

We developed an atom–cluster synergistic strategy to prepare Fe single atoms and nanoclusters coexisting nanozyme (Fe_SA+NC_), which preferred to decompose H_2_O_2_ through CAT-like pathway while suppressing POD-like route. Experiments and DFT calculations revealed that introducing Fe nanoclusters resulted in a reduced local symmetry of FeN_4_ sites, electron redistribution, and a higher spin state. These modulations optimized the adsorption energy of OH and H, suppressed ^•^OH release, and facilitated the O–O bond cleavage, thereby improving selectivity toward CAT-like activity. Importantly, this CAT-preferred and POD-suppressed behavior is desirable for anti-inflammatory therapy. Using a rat RA model as a proof-of-concept, we demonstrated the therapeutic potential of Fe_SA+NC_ as an anti-inflammatory agent. Moreover, Fe_SA+NC_ showed promise as an adjunct to stem cell therapy by mitigating oxidative stress in the local microenvironment. Overall, this work highlights a strategy to tune reaction selectivity among multiple enzyme-like activities in nanozymes. This design principle is expected to be extendable to other SAzymes systems, offering a general strategy for developing controllable and efficient nanozyme-based therapeutics.

## Acknowledgments

This work was supported by the National Natural Science Foundation of China, W2512074, 22374071 (to H.W.), and 22303020 (to Z.W.). the Key Program of Nanozyme Laboratory in Zhongyuan, NLZ-KP2024NIC06 (to H.W.), the Jiangsu Provincial Key R&D Program, BE2022836 (to H.W.), the State Key Laboratory of Analytical Chemistry for Life Science, 5431ZZXM2501 (to H.W.), Fundamental Research Funds for the Central Universities, 2025300292 (to H.W.), the Open Funds of NMPA Key Laboratory for Biomedical Optics, 20240001 (to H.W.), PAPD Program, the International Expansion and Enhancement Program by Nanjing University International Affairs Office, and Yachen Foundation of Nanjing University.

## References

[1] Di Matteo, A.; Bathon, J. M.; Emery, P. Rheumatoid Arthritis. The Lancet 2023, 402, 2019–2033.

[2] Firestein, G. S. Evolving Concepts of Rheumatoid Arthritis. Nature 2003, 423, 356–361.

[3] Schieber, M.; Chandel, Navdeep S. ROS Function in Redox Signaling and Oxidative Stress. Current Biology 2014, 24, R453–R462.

[4] Forman, H. J.; Zhang, H. Targeting Oxidative Stress in Disease: Promise and Limitations of Antioxidant Therapy. Nature Reviews Drug Discovery 2021, 20, 689–709.

[5] Li, B.; Ming, H.; Qin, S.; Nice, E. C.; Dong, J.; Du, Z.; Huang, C. Redox Regulation: Mechanisms, Biology and Therapeutic Targets in Diseases. Signal Transduction and Targeted Therapy 2025, 10, 72.

[6] Azadmanesh, J.; Slobodnik, K.; Struble, L. R.; Lutz, W. E.; Coates, L.; Weiss, K. L.; Myles, D. A. A.; Kroll, T.; Borgstahl, G. E. O. Revealing the Atomic and Electronic Mechanism of Human Manganese Superoxide Dismutase Product Inhibition. Nature Communications 2024, 15, 5973.

[7] Nandi, A.; Yan, L.-J.; Jana, C. K.; Das, N. Role of Catalase in Oxidative Stress- and Age-Associated Degenerative Diseases. Oxidative Medicine and Cellular Longevity 2019, 2019, 9613090.

[8] Wu, J.; Wang, X.; Wang, Q.; Lou, Z.; Li, S.; Zhu, Y.; Qin, L.; Wei, H. Nanomaterials with Enzyme-Like Characteristics (nanozymes): Next-Generation Artificial Enzymes (II). Chemical Society Reviews 2019, 48, 1004–1076.

[9] Liu, Y.; Cheng, Y.; Zhang, H.; Zhou, M.; Yu, Y.; Lin, S.; Jiang, B.; Zhao, X.; Miao, L.; Wei, C.-W.; Liu, Q.; Lin, Y.-W.; Du, Y.; Butch, C. J.; Wei, H. Integrated Cascade Nanozyme Catalyzes in vivo ROS Scavenging for Anti-inflammatory Therapy. Science Advances 2020, 6, eabb2695.

[10] Yuan, B.; Tan, Z.; Guo, Q.; Shen, X.; Zhao, C.; Chen, J. L.; Peng, Y.-K. Regulating the H_2_O_2_ Activation Pathway on a Well-Defined CeO_2_ Nanozyme Allows the Entire Steering of Its Specificity between Associated Enzymatic Reactions. ACS Nano 2023, 17, 17383–17393.

[11] Ji, S.; Jiang, B.; Hao, H.; Chen, Y.; Dong, J.; Mao, Y.; Zhang, Z.; Gao, R.; Chen, W.; Zhang, R.; Liang, Q.; Li, H.; Liu, S.; Wang, Y.; Zhang, Q.; Gu, L.; Duan, D.; Liang, M.; Wang, D.; Yan, X.; Li, Y. Matching the Kinetics of Natural Enzymes with a Single-Atom Iron Nanozyme. Nature Catalysis 2021, 4, 407–417.

[12] Huang, L.; Chen, J.; Gan, L.; Wang, J.; Dong, S. Single-Atom Nanozymes. Science Advances 2019, 5, eaav5490.

[13] Wu, J.; Zhu, X.; Li, Q.; Fu, Q.; Wang, B.; Li, B.; Wang, S.; Chang, Q.; Xiang, H.; Ye, C.; Li, Q.; Huang, L.; Liang, Y.; Wang, D.; Zhao, Y.; Li, Y. Enhancing Radiation-Resistance and Peroxidase-like Activity of Single-Atom Copper Nanozyme via Local Coordination Manipulation. Nature Communications 2024, 15, 6174.

[14] Wei, S.; Ma, W.; Sun, M.; Xiang, P.; Tian, Z.; Mao, L.; Gao, L.; Li, Y. Atom-Pair Engineering of Single-Atom Nanozyme for Boosting Peroxidase-Like Activity. Nature Communications 2024, 15, 6888.

[15] Chen, Y.; Jiang, B.; Hao, H.; Li, H.; Qiu, C.; Liang, X.; Qu, Q.; Zhang, Z.; Gao, R.; Duan, D.; Ji, S.; Wang, D.; Liang, M. Atomic-Level Regulation of Cobalt Single-Atom Nanozymes: Engineering High-Efficiency Catalase Mimics. Angewandte Chemie International Edition 2023, 62, e202301879.

[16] Shang, H.; Zhou, X.; Dong, J.; Li, A.; Zhao, X.; Liu, Q.; Lin, Y.; Pei, J.; Li, Z.; Jiang, Z.; Zhou, D.; Zheng, L.; Wang, Y.; Zhou, J.; Yang, Z.; Cao, R.; Sarangi, R.; Sun, T.; Yang, X.; Zheng, X.; Yan, W.; Zhuang, Z.; Li, J.; Chen, W.; Wang, D.; Zhang, J.; Li, Y. Engineering Unsymmetrically Coordinated Cu–S_1_N_3_ Single Atom Sites with Enhanced Oxygen Reduction Activity. Nature Communications 2020, 11, 3049.

[17] Wei, S.; Sun, M.; Huang, J.; Chen, Z.; Wang, X.; Gao, L.; Zhang, J. Axial Chlorination Engineering of Single-Atom Nanozyme: Fe–N_4_Cl Catalytic Sites for Efficient Peroxidase-Mimicking. Journal of the American Chemical Society 2024, 146, 33239–33248.

[18] Liu, Y.; Wang, B.; Zhu, J.; Xu, X.; Zhou, B.; Yang, Y. Single-Atom Nanozyme with Asymmetric Electron Distribution for Tumor Catalytic Therapy by Disrupting Tumor Redox and Energy Metabolism Homeostasis. Advanced Materials 2023, 35, 2208512.

[19] Li, Z.; Liu, F.; Chen, C.; Jiang, Y.; Ni, P.; Song, N.; Hu, Y.; Xi, S.; Liang, M.; Lu, Y. Regulating the N Coordination Environment of Co Single-Atom Nanozymes for Highly Efficient Oxidase Mimics. Nano Letters 2023, 23, 1505–1513.

[20] Yang, Q.; Liu, J.; Cai, W.; Liang, X.; Zhuang, Z.; Liao, T.; Zhang, F.; Hu, W.; Liu, P.; Fan, S.; Yu, W.; Jiang, B.; Li, C.; Wang, D.; Xu, Z. Non-Heme Iron Single-Atom Nanozymes as Peroxidase Mimics for Tumor Catalytic Therapy. Nano Letters 2023, 23, 8585–8592.

[21] Wang, L.; Liu, Z.; Yao, L.; Liu, S.; Wang, Q.; Qu, H.; Wu, Y.; Mao, Y.; Zheng, L. A Bioinspired Single-Atom Fe Nanozyme with Excellent Laccase-Like Activity for Efficient Aflatoxin B1 Removal. Small 2024, 20, 2400629.

[22] Liu, W.; Shi, E.; Wu, H.; Liang, Y.; Chen, M.; Zhang, H.; Zhang, R.; Li, X.; Wang, Y.; Zhang, L. Spatially Axial Boron Coordinated Single-Atom Nanozymes with Boosted Multi-Enzymatic Performances for Periodontitis Treatment. Advanced Functional Materials 2024, 34, 2403386.

[23] Zhong, S.; Zhang, Z.; Wang, Z.; Zhao, Q.; Chen, W.; Chen, G.; Jiang, Z.; Cai, Q.; Gong, L.; Lai, Y.; Wang, D.; Li, L. Synergizing Catalysis with Post-catalysis Pseudo-Iron Release by Building Dynamic Catalytic Active Sites in Diatomic Nanozymes for Boosting Cancer Therapy. Journal of the American Chemical Society 2025, 147, 15814–15826.

[24] Ye, J.; Li, C.; Xu, J.; Liu, S.; Qu, J.; Wang, Q.; Cao, J.; Zhao, Y.; Li, C.; Yang, P. Engineered Nanozymes with Asymmetric Mn–O–Ce Sites for Intratumorally Leveraged Multimode Therapy. Advanced Materials 2025, 37, 2419673.

[25] Zhang, H.; Chen, H.-C.; Feizpoor, S.; Li, L.; Zhang, X.; Xu, X.; Zhuang, Z.; Li, Z.; Hu, W.; Snyders, R.; Wang, D.; Wang, C. Tailoring Oxygen Reduction Reaction Kinetics of Fe–N–C Catalyst via Spin Manipulation for Efficient Zinc-Air Batteries. Advanced Materials 2024, 36, 2400523.

[26] Sun, Z.; Shi, W.; Smith, L. R.; Dummer, N. F.; Qi, H.; Sun, Z.; Hutchings, G. J. Concerted Catalysis of Single Atom and Nanocluster Enhances Bio-ethanol Activation and Dehydrogenation. Nature Communications 2025, 16, 3935.

[27] Jiang, W.-J.; Gu, L.; Li, L.; Zhang, Y.; Zhang, X.; Zhang, L.-J.; Wang, J.-Q.; Hu, J.-S.; Wei, Z.; Wan, L.-J. Understanding the High Activity of Fe–N–C Electrocatalysts in Oxygen Reduction: Fe/Fe_3_C Nanoparticles Boost the Activity of Fe–N_x_. Journal of the American Chemical Society 2016, 138, 3570–3578.

[28] Cui, X.; Gao, L.; Lei, S.; Liang, S.; Zhang, J.; Sewell, C. D.; Xue, W.; Liu, Q.; Lin, Z.; Yang, Y. Simultaneously Crafting Single-Atomic Fe Sites and Graphitic Layer-Wrapped Fe_3_C Nanoparticles Encapsulated within Mesoporous Carbon Tubes for Oxygen Reduction. Advanced Functional Materials 2021, 31, 2009197.

[29] Yuan, L.; Liu, B.; Shen, L.; Dai, Y.; Li, Q.; Liu, C.; Gong, W.; Sui, X.; Wang, Z. d-Orbital Electron Delocalization Realized by Axial Fe_4_C Atomic Clusters Delivers High-Performance Fe–N–C Catalysts for Oxygen Reduction Reaction. Advanced Materials 2023, 35, 2305945.

[30] Mo, F.; Song, C.; Zhou, Q.; Xue, W.; Ouyang, S.; Wang, Q.; Hou, Z.; Wang, S.; Wang, J. The Optimized Fenton-Like Activity of Fe Single-Atom Sites by Fe Atomic Clusters-Mediated Electronic Configuration Modulation. Proceedings of the National Academy of Sciences 2023, 120, e2300281120.

[31] Zhang, J.; Pan, Y.; Feng, D.; Cui, L.; Zhao, S.; Hu, J.; Wang, S.; Qin, Y. Mechanistic Insight into the Synergy between Platinum Single Atom and Cluster Dual Active Sites Boosting Photocatalytic Hydrogen Evolution. Advanced Materials 2023, 35, 2300902.

[32] Yang, X.; Wang, Y.; Wang, X.; Mei, B.; Luo, E.; Li, Y.; Meng, Q.; Jin, Z.; Jiang, Z.; Liu, C.; Ge, J.; Xing, W. CO-Tolerant PEMFC Anodes Enabled by Synergistic Catalysis between Iridium Single-Atom Sites and Nanoparticles. Angewandte Chemie International Edition 2021, 60, 26177–26183.

[33] Qiao, Y.; Yuan, P.; Hu, Y.; Zhang, J.; Mu, S.; Zhou, J.; Li, H.; Xia, H.; He, J.; Xu, Q. Sulfuration of an Fe–N–C Catalyst Containing FeC/Fe Species to Enhance the Catalysis of Oxygen Reduction in Acidic Media and for Use in Flexible Zn-Air Batteries. Advanced Materials 2018, 30, 1804504.

[34] Vorontsov, A. V.; Tsybulya, S. V. Influence of Nanoparticles Size on XRD Patterns for Small Monodisperse Nanoparticles of Cu^0^ and TiO_2_ Anatase. Industrial & Engineering Chemistry Research 2018, 57, 2526–2536.

[35] Li, J.; Chen, M.; Cullen, D. A.; Hwang, S.; Wang, M.; Li, B.; Liu, K.; Karakalos, S.; Lucero, M.; Zhang, H.; Lei, C.; Xu, H.; Sterbinsky, G. E.; Feng, Z.; Su, D.; More, K. L.; Wang, G.; Wang, Z.; Wu, G. Atomically Dispersed Manganese Catalysts for Oxygen Reduction in Proton-exchange Membrane Fuel Cells. Nature Catalysis 2018, 1, 935–945.

[36] Sheng, Z.; Shao, L.; Chen, J.; Bao, W.; Wang, F.; Xia, X. Catalyst-Free Synthesis of Nitrogen-Doped Graphene via Thermal Annealing Graphite Oxide with Melamine and Its Excellent Electrocatalysis. ACS Nano 2011, 5, 4350–4358.

[37] Lazar, P.; Mach, R.; Otyepka, M. Spectroscopic Fingerprints of Graphitic, Pyrrolic, Pyridinic, and Chemisorbed Nitrogen in N-Doped Graphene. The Journal of Physical Chemistry C 2019, 123, 10695–10702.

[38] Roh, J.; Cho, A.; Kim, S.; Lee, K.-S.; Shin, J.; Choi, J. S.; Bak, J.; Lee, S.; Song, D.; Kim, E.-J.; Lee, C.; Uhm, Y. R.; Cho, Y.-H.; Han, J. W.; Cho, E. Transformation of the Active Moiety in Phosphorus-Doped Fe–N–C for Highly Efficient Oxygen Reduction Reaction. ACS Catalysis 2023, 13, 9427–9441.

[39] Li, Z.; Wu, D.; Wang, Q.; Zhang, Q.; Xu, P.; Liu, F.; Xi, S.; Ma, D.; Lu, Y.; Jiang, L.; Zhang, Z. Bioinspired Homonuclear Diatomic Iron Active Site Regulation for Efficient Antifouling Osmotic Energy Conversion. Advanced Materials 2024, 36, 2408364.

[40] Li, Q.; Liu, H.; Zhang, L.; Chen, H.; Zhu, H.; Wu, Y.; Xu, M.; Bao, S. Highly Efficient Fe–N–C Oxygen Reduction Electrocatalyst Engineered by Sintering Atmosphere. Journal of Power Sources 2020, 449, 227497.

[41] Zeng, Y.; Li, C.; Li, B.; Liang, J.; Zachman, M. J.; Cullen, D. A.; Hermann, R. P.; Alp, E. E.; Lavina, B.; Karakalos, S.; Lucero, M.; Zhang, B.; Wang, M.; Feng, Z.; Wang, G.; Xie, J.; Myers, D. J.; Dodelet, J.-P.; Wu, G. Tuning the Thermal Activation Atmosphere Breaks the Activity-Stability Trade-off of Fe–N–C Oxygen Reduction Fuel Cell Catalysts. Nature Catalysis 2023, 6, 1215–1227.

[42] Ouyang, W.; Zeng, D.; Yu, X.; Xie, F.; Zhang, W.; Chen, J.; Yan, J.; Xie, F.; Wang, L.; Meng, H.; Yuan, D. Exploring the Active Sites of Nitrogen-Doped Graphene as Catalysts for the Oxygen Reduction Reaction. International Journal of Hydrogen Energy 2014, 39, 15996–16005.

[43] Ning, X.; Li, Y.; Ming, J.; Wang, Q.; Wang, H.; Cao, Y.; Peng, F.; Yang, Y.; Yu, H. Electronic Synergism of Pyridinic- and Graphitic-Nitrogen on N-Doped Carbons for the Oxygen Reduction Reaction. Chemical Science 2019, 10, 1589–1596.

[44] Xu, Y.; Mo, Y.; Tian, J.; Wang, P.; Yu, H.; Yu, J. The Synergistic Effect of Graphitic N and Pyrrolic N for the Enhanced Photocatalytic Performance of Nitrogen-doped Graphene/TiO_2_ Nanocomposites. Applied Catalysis B: Environmental 2016, 181, 810–817.

[45] Yang, G.; Cai, H.; Zhang, N.; Wang, B.; Liang, C.; Zhang, S.; Yang, Z.; Yang, S. Regulation of d-Orbital Electron in Fe–N_4_ by High-Entropy Atomic Clusters for Highly Active and Durable Oxygen Reduction Reaction. Advanced Functional Materials 2024, 34, 2407775.

[46] Westre, T. E.; Kennepohl, P.; DeWitt, J. G.; Hedman, B.; Hodgson, K. O.; Solomon, E. I. A Multiplet Analysis of Fe K-Edge 1s → 3d Pre-Edge Features of Iron Complexes. Journal of the American Chemical Society 1997, 119, 6297–6314.

[47] Lu, X.; Kuai, L.; Huang, F.; Jiang, J.; Song, J.; Liu, Y.; Chen, S.; Mao, L.; Peng, W.; Luo, Y.; Li, Y.; Dong, H.; Li, B.; Shi, J. Single-atom catalysts-based catalytic ROS clearance for efficient psoriasis treatment and relapse prevention via restoring ESR1. Nature Communications 2023, 14, 6767.

[48] Zhang, R.; Xue, B.; Tao, Y.; Zhao, H.; Zhang, Z.; Wang, X.; Zhou, X.; Jiang, B.; Yang, Z.; Yan, X.; Fan, K. Edge-Site Engineering of Defective Fe-N_4_ Nanozymes with Boosted Catalase-Like Performance for Retinal Vasculopathies. Advanced Materials 2022, 34, 2205324.

[49] Sies, H.; Jones, D. P. Reactive Oxygen Species (ROS) as Pleiotropic Physiological Signalling Agents. Nature Reviews Molecular Cell Biology 2020, 21, 363–383.

[50] Sies, H. Hydrogen Peroxide as a Central Redox Signaling Molecule in Physiological Oxidative Stress: Oxidative Eustress. Redox Biology 2017, 11, 613–619.

[51] Zhang, R.; Yan, X.; Gao, L.; Fan, K. Nanozymes Expanding the Boundaries of Biocatalysis. Nature Communications 2025, 16, 6817.

[52] Kim, H. S.; Lee, S.; Lee, D. Y. Aurozyme: A Revolutionary Nanozyme in Colitis, Switching Peroxidase-Like to Catalase-Like Activity. Small 2023, 19, 2302331.

[53] Cursi, L.; Mirra, G.; Boselli, L.; Pompa, P. P. Metrology of Platinum Nanozymes: Mechanistic Insights and Analytical Issues. Advanced Functional Materials 2024, 34, 2315587.

[54] Zang, Y.; Liu, Y.; Lu, R.; Yang, Q.; Wang, B.; Zhang, M.; Mao, Y.; Wang, Z.; Lum, Y. Tuning Transition Metal 3d Spin state on Single-atom Catalysts for Selective Electrochemical CO_2_ Reduction. Advanced Materials 2025, 37, 2417034.

[55] Li, Z.; Zhuang, Z.; Lv, F.; Zhu, H.; Zhou, L.; Luo, M.; Zhu, J.; Lang, Z.; Feng, S.; Chen, W.; Mai, L.; Guo, S. The Marriage of the FeN_4_ Moiety and MXene Boosts Oxygen Reduction Catalysis: Fe 3d Electron Delocalization Matters. Advanced Materials 2018, 30, 1803220.

[56] Shen, X.; Wang, Z.; Gao, X.; Zhao, Y. Density Functional Theory-Based Method to Predict the Activities of Nanomaterials as Peroxidase Mimics. ACS Catalysis 2020, 10, 12657–12665.

[57] Gao, X. J.; Yan, J.; Zheng, J.; Zhong, S.; Gao, X. Clear-Box Machine Learning for Virtual Screening of 2D Nanozymes to Target Tumor Hydrogen Peroxide. Advanced Healthcare Materials 2023, 12, 2202925.

[58] Pan, L.; Hu, X.; Yang, H.; Chen, L.; Tang, H.; Luo, S.; Hu, H.; Zhang, J. Nanotherapy-Engineered Stem Cells Alleviate Myocardial Reperfusion Injury by Normalizing the Pathological Microenvironment and Promoting Cardiac Repair. Advanced Functional Materials 2025, 35, 2417338.

[59] Le, W.; Sun, Z.; Li, T.; Cao, H.; Yang, C.; Mei, T.; Zhang, L.; Wang, Y.; Jia, W.; Sun, W.; Hu, Y.; Liu, Z. Antioxidant Nanozyme-Engineered Mesenchymal Stem Cells for In Vivo MRI Tracking and Synergistic Therapy of Myocardial Infarction. Advanced Functional Materials 2024, 34, 2314328.

[60] Zhu, H.; Wu, X.; Liu, R.; Zhao, Y.; Sun, L. ECM-Inspired Hydrogels with ADSCs Encapsulation for Rheumatoid Arthritis Treatment. Advanced Science 2023, 10, 2206253.

[61] You, D. G.; Lim, G. T.; Kwon, S.; Um, W.; Oh, B. H.; Song, S. H.; Lee, J.; Jo, D.-G.; Cho, Y. W.; Park, J. H. Metabolically Engineered Stem Cell-Derived Exosomes to Regulate Macrophage Heterogeneity in Rheumatoid Arthritis. Science Advances 2021, 7, eabe0083.

[62] Zhao, Y.; Song, S.; Wang, D.; Liu, H.; Zhang, J.; Li, Z.; Wang, J.; Ren, X.; Zhao, Y. Nanozyme-Reinforced Hydrogel as a H_2_O_2_-Driven Oxygenerator for Enhancing Prosthetic Interface Osseointegration in Rheumatoid Arthritis Therapy. Nature Communications 2022, 13, 6758.

[63] Yang, B.; Yao, H.; Yang, J.; Chen, C.; Shi, J. Construction of a Two-Dimensional Artificial Antioxidase for Nanocatalytic Rheumatoid Arthritis Treatment. Nature Communications 2022, 13, 1988.

[64] Ma, Y.; Lu, Z.; Jia, B.; Shi, Y.; Dong, J.; Jiang, S.; Li, Z. DNA Origami as a Nanomedicine for Targeted Rheumatoid Arthritis Therapy through Reactive Oxygen Species and Nitric Oxide Scavenging. ACS Nano 2022, 16, 12520–12531.

