## Supporting Information for "Spin-State Modulation by Atom–Cluster Synergy Steers H2O2 Conversion toward a Catalase-like Decomposition Pathway for Anti-Inflammatory Therapy"

#### Experimental Section

##### Chemicals and reagents

$Zn(NO_3)_2 \cdot 6H_2O$, iron(III) acetylacetonate ($Fe(acac)_3$), and 3,3′,5,5′-tetramethylbenzidine (TMB) were purchased from Aladdin Chemical Co., Ltd. (Shanghai, China). Hydrogen peroxide ($H_2O_2$, 30 wt% in water) and methanol were purchased from Sinopharm Chemical Reagent Co., Ltd. (Shanghai, China). 2-methylimidazole (2-MeIM; 99%) was purchased from Bidepharm (Shanghai, China). SOD Assay Kit-WST was purchased from DOJINDO Laboratories (Kumamoto, Japan). 5,5-Dimethyl-1-pyrroline N-oxide (DMPO), Cell Counting Kit-8 (CCK-8), and 2',7'-dichlorodihydrofluorescein diacetate (DCFH-DA) were purchased from MedChemExpress (New Jersey, USA). APC anti-mouse CD86 and PE anti-mouse CD206 were purchased from Biolegend (California, USA). All chemicals were purchased from commercial sources and used directly without further purification. Ultrapure water (18 MΩ•cm) was obtained from a Milli-Q purification system.

##### Materials characterization

Powder X-ray diffraction (XRD) patterns of nanoparticles were measured by using an Ultima IV diffractometer (Rigaku, Japan). X-ray photoelectron spectroscopy (XPS) spectra were collected by using a PHI 5000 Versa Probe XPS microscope (Ulvac-Phi, Japan). Transmission electron microscopy (TEM) and scanning electron microscopy (SEM) images were obtained from JEM-1400Flash (JEOL, Japan) and Ultra 55 microscope (Zeiss, Germany), respectively. High-angle annular dark-field scanning transmission electron microscopy (HAADF-STEM) images were captured using an FEI Themis Z instrument equipped with a probe corrector and monochromator operating at 200 kV. Zeta potential and particle size were measured using a Malvern Zetasizer Nano ZS90. Raman spectra were acquired from an inVia-Reflex spectrometer (Renishaw, England). The Brunauer–Emmett–Teller (BET) surface area and pore size were determined by using a Kubo X1000 instrument (Builder, China). Inductively coupled plasma (ICP) was performed by using an ICAP7400 ICP-OES instrument (Thermo Fisher Scientific, USA). Electron paramagnetic resonance (EPR) spectra were recorded on an A300 spectrometer (Bruker, USA). Ultraviolet–visible absorption spectra were measured by using a SpectraMax M2e microplate reader (Molecular Devices, USA) or a UV-3600 Plus UV–vis spectrophotometer (Shimadzu, Japan). Dissolved oxygen experiment was performed by using an InLab OptiOx detector (Mettler Toledo, Switzerland). Flow cytometry experiment was carried out by using a Cytoflex instrument (Beckman, USA). The three-dimensional scan of the tissue of SD rat was obtained via a vivaCT 80 instrument (Scanco Medical AG, Switzerland). The staining images of cell and tissue were photographed by using a DMi8 fluorescence microscope (Leica, Germany).

##### Measurements and analysis of XAS

The X-ray absorption fine structure spectra (XAFS) were collected at BL-14W1 of Shanghai Synchrotron Radiation Facility (SSRF), China. The data collection was carried out in transmission mode using ionization chamber for Fe foil, and in fluorescence excitation mode using a Lytle detector for $Fe_{SA}$ and $Fe_{SA+NC}$. All spectra were collected in ambient conditions. The XAFS data were processed according to the standard procedures using the Athena module implemented in the IFEFFIT software packages. The extended X-ray absorption fine structure (EXAFS) spectra were obtained by subtracting the post-edge background from the overall absorption and then normalizing with respect to the edge-jump step. Subsequently, the $\chi(k)$ data of were Fourier transformed into real

(R) space using a hanning windows ($dk$ = 1.0 Å$^{-1}$) to separate the EXAFS contributions from different coordination shells. To obtain the quantitative structural parameters around central atoms, least-squares curve parameter fitting was performed using the ARTEMIS module of IFEFFIT software packages.

**Synthesis of Fe/ZIF-8**

3.94 g of 2-methylimidazole was dissolved completely in 60 mL methanol solution with stirring. 3.57 g of $Zn(NO_3)\cdot 6H_2O$ and 211 mg of $Fe(acac)_3$ were dissolved in 90 mL methanol solution under ultrasound for 10 minutes. Then, the two solutions were mixed and stirred vigorously for 6 h at room temperature. After that, the precipitant was collected by centrifuging, washed with methanol four times and dried at 60 °C in vacuum for overnight. ZIF-8 was synthesized by similar procedure, except that $Fe(acac)_3$ was not added.

**Synthesis of $Fe_{SA+NC}$ and $Fe_{SA}$**

After grinding, 200 mg of Fe/ZIF-8 powder was put into ceramic boat and transferred to a tube furnace. Next, the powder was pyrolyzed under 900 ℃ in a 10% $H_2$ in Ar atmosphere for 3 h to synthesize $Fe_{SA+NC}$. For comparison, the same precursor was pyrolyzed at 900 ℃ in Ar to synthesize $Fe_{SA}$. The heating rate was 5 °C min$^{-1}$.

**Synthesis of $Fe_{NC}$**

ZIF8-derived N-doped carbon (CN) was first loaded with Fe species by an incipient wetness impregnation method. Typically, 50 mg of CN was impregnated with an aqueous $Fe(NO_3)_3\cdot 9H_2O$ solution (10 mg mL$^{-1}$) added dropwise under continuous stirring until the support was just saturated. The resulting wet solid was thoroughly mixed and then dried in an oven at 60 °C. Then, the dried precursor was subsequently subjected to programmed reduction in flowing 10% $H_2$/Ar atmosphere. The sample was heated from room temperature to 200 °C and kept at this temperature for 1 h, then further heated to 350 °C and maintained for another 1 h. Afterwards, the temperature was increased to 600 °C and held for 30 min. The heating rate was 2 °C min$^{-1}$.

**Sulfuric acid washing and KSCN poisoning experiment**

To remove Fe nanoclusters, $Fe_{SA+NC}$ was dispersed in 20 mL of 0.5 M $H_2SO_4$ and stirred at 80 °C for 12 h. After cooling to room temperature, the solid was collected by centrifuge, thoroughly washed with deionized water until the supernatant became neutral (pH ≈ 7), and then dried at 60 °C in vacuum for overnight.

For the poisoning experiments, $Fe_{SA+NC}$ was incubated with KSCN at room temperature for 24 h under gentle stirring. The solid was then collected by centrifugation, washed once with fresh buffer to remove excess free KSCN and finally re-suspended in buffer for subsequent activity measurements.

**CAT-like activity assay**

Two methods were used to compare the CAT-like activity of $Fe_{SA+NC}$ and $Fe_{SA}$. Firstly, $H_2O_2$ (5 mM) and nanozymes (5 μg mL$^{-1}$) were mixed in 0.1 M PBS buffer (pH 7.4). The oxygen solubility (mg L$^{-1}$) was continuously monitored every 5 s by the dissolved oxygen meter. Secondly, $H_2O_2$ (20 mM) and nanozymes with different concentrations (5, 10, 20, 40 μg mL$^{-1}$) were added to 0.1 M PBS buffer (pH 7.4) within 1 mL quartz cuvette. The absorbance changes of the reaction solution at 240 nm were recorded using a UV spectrophotometer immediately. Then, the specific CAT-like activity of nanozymes was calculated according to the equation (1):

$$Catalase\ (U\ mg^{-1}) = \frac{\Delta A_{240\,nm}\cdot V\cdot 10^6}{\Delta t\cdot \varepsilon\cdot d\cdot m} \quad (1)$$

$\Delta A_{240\ nm}$ was the absorbance change; V was the volume of the reaction solution (0.001 L); Δt was the reaction time (s); ε was the extinction coefficient of $H_2O_2$ (43.6 $M^{-1}$ $cm^{-1}$) at 240 nm; d was the optical path length (1 cm); m was the mass of nanozyme (mg).

The pH dependence of nanozymes' CAT-like activity was also measured by the above methods.

**Kinetics study of CAT-like activity**

Typically, 20 μL of nanozymes (1 mg $mL^{-1}$) were added to 0.1 M PBS (pH 7.4), followed by the introduction of 20 μL of $H_2O_2$ with different final concentrations (1, 5, 10, 15, 20 mM). The absorbance changes of the reaction solution at 240 nm were recorded using a UV spectrophotometer immediately. The initial velocities were calculated based on the Beer-Lambert Law. Then the Michaelis-Menten curve was produced by plotting the calculated initial rate against the substrate concentrations. The maximal velocities ($V_{max}$) and Michaelis constants ($K_m$) were obtained based on the nonlinear least squares regression analysis of the Michaelis-Menten curve.

**SOD-like activity assay**

The SOD-like activity of nanozymes were tested by using SOD Assay Kit (S311, DOJINDO Molecular Technologies) according to the provided protocol. In brief, 200 μL of diluting WST-1 working solution and 20 μL of enzyme working solution were added to 20 μL of nanozyme solution in 96-well plates. The final concentration of nanozymes were 2.5, 5, 10, 15, 20 μg $mL^{-1}$. After thoroughly mixing and incubation at 37 °C for 20 minutes, the absorbance at 450 nm was measured using a microplate reader. The specific SOD activity of nanozymes was calculated according to the equation (2):

$$SOD(U\ mg^{-1}) = \frac{1}{IC_{50} \cdot V} \tag{2}$$

$IC_{50}$ was the concentration of nanozyme that could remove 50 percent $O_2^{\bullet-}$; V was the volume of the reaction solution (240 μL).

**POD-like activity assay**

Typically, 20 μL of nanozymes (1 mg $mL^{-1}$) were added to 0.2 M HAc-NaAc buffer (pH 4.5), followed by the introduction of $H_2O_2$ and TMB with different final concentrations. Colorimetric reactions were recorded in time-scan mode by measuring the absorbance at 652 nm using a UV spectrophotometer immediately. The initial velocities were calculated based on the Beer-Lambert Law. Then the Michaelis-Menten curve was produced by plotting the calculated initial rate against the substrate concentrations. The maximal velocities ($V_{max}$) and Michaelis constants ($K_m$) were obtained based on the nonlinear least squares regression analysis of the Michaelis-Menten curve.

In addition to the specific POD-like activity of nanozymes, TMB (1 mM) and a much higher concentration of $H_2O_2$ (1 M) were used to measurement. The specific POD-like activity was calculated according to the equation (3):

$$Peroxidase\ (U\ mg^{-1}) = \frac{\Delta A_{652\ nm} \cdot V \cdot 10^6}{\Delta t \cdot \varepsilon \cdot d \cdot m} \tag{3}$$

$\Delta A_{652\ nm}$ was the absorbance change; V was the volume of the reaction solution (0.001 L); Δt was the reaction time (s); ε was the extinction coefficient of TMB (39000 $M^{-1}$ $cm^{-1}$) at 652 nm; d was the optical path length (1 cm); m was the mass of nanozyme (mg).

**OXD-like activity assay**

The OXD-like activity was measured under the same condition as in the POD-like activity assay but without the addition of $H_2O_2$. In addition to the specific OXD-like activity of nanozymes was calculated according to the equation (4):

$$Oxidase\ (U\ mg^{-1}) = \frac{\Delta A_{652\ nm} \cdot V \cdot 10^6}{\Delta t \cdot \varepsilon \cdot d \cdot m} \tag{4}$$

$\Delta A_{652\ nm}$ was the absorbance change; V was the volume of the reaction solution (0.001 L); Δt was the reaction time (s); ε was the extinction coefficient of TMB (39000 $M^{-1}$ $cm^{-1}$) at 652 nm; d was the optical path length (1 cm); m was the mass of nanozyme (mg).

**•OH scavenging**

Typically, $Fe^{2+}$ (1 mM) and $H_2O_2$ (1 mM) were mixed in 0.1 M PBS buffer (pH 7.4) to generate •OH. Then, DMPO (10 mM) and nanozymes were added into above solution. After 90s, the reaction solution was drawn into a quartz capillary and placed in a glass tube for EPR analysis.

**Computational details**

Density functional theory (DFT) calculations were performed using the projector augmented-wave (PAW) method, as implemented in the Vienna ab initio Simulation Package (VASP). [1] The generalized gradient approximation proposed by Perdew, Burke, and Ernzerhof was selected for the exchange-correlation potential. [2] The long range van der Waals interaction was described by the DFT-D3(BJ) approach. [3] The cut-off energy for plane wave was set to 400 eV. The energy criterion was set to $10^{-5}$ eV in iterative solution of the Kohn-Sham equation. Force criterion was set to 0.02 eV $Å^{-1}$. The Brillouin zone integration was performed using a 3×3×1 k-mesh for $FeN_4$ and $Fe_4N_6$, and a 5×5×1 k-mesh for density of states (DOS) calculations. A vacuum layer of 15 Å was applied to avoid interactions between periodic images. The adsorption energy ($E_{ads}$) was calculated according to the equation (5):

$$\Delta E_{ads} = E_{complex} - E_{model} - E_{mol} \tag{5}$$

$E_{complex}$, $E_{model}$, and $E_{mol}$ represented the energy of complex, pure model and chemical group, respectively.

**Cell culture**

RAW 264.7 cells were obtained from the cell bank of the Chinese Academy of Sciences (Shanghai, China). RAW264.7 cells were incubated in High glucose-Dulbecco's modified eagle medium (DMEM) containing 10% fetal bovine serum (FBS) and 1% penicillin/streptomycin in a humidified 5% $CO_2$ atmosphere at 37 °C. BMSCs were isolated from the bone marrow of 2-week-old Sprague-Dawley (SD) rats. The conditions of culture were consistent with RAW 264.7 cells except the medium was DMEM/F12.

**Cell viability assay**

Briefly, RAW264.7 cells and BMSCs were plated into 96-well plates at 1 × $10^4$ cells per well. The medium containing nanozymes were refreshed after overnight incubation for adherence. Following incubation for another 24 h, CCK-8 kits were used to evaluate the cell viability.

**Intracellular ROS scavenging**

Briefly, RAW264.7 cells were plated into 12-well plates at 1 × $10^5$ cells per well and allowed to adhere after incubation overnight. Different amounts of Nanozymes were added into wells and incubated for another 2 h. Then, the medium was replaced with fresh medium containing $H_2O_2$ (100 μM) for oxidative stimulation, lasting 30 minutes. Following that, intracellular ROS level was measured by fluorescence intensities stained with DCHF-DA probe.

**Cell protective function assay**

Briefly, BMSCs were plated into 12-well plates at 1×$10^5$ cells per well and allowed to adhere after incubation overnight. After cultured with nanozyme for 24 h, the cells were treated with $H_2O_2$ (100 μM) for another 24 h. Then, Calcein/PI Cell Viability/Cytotoxicity Assay Kit was used to evaluate the cytoprotective ability of nanozyme.

**Macrophage reprograming**

Typically, RAW264.7 cells were plated into 12-well plates at $1 \times 10^5$ cells per well and allowed to adhere after incubation overnight. Then, nanozyme, BMSC and Fe/BMSC were co-cultured with LPS (100 ng $mL^{-1}$) stimulated macrophages for 24 h. Subsequently, the macrophages were stained with anti-APC-CD86 and anti-PE-CD206 for detection by flow cytometry.

**Animal experiment**

All animal experiments were approved by the Institutional Animal Care and Use Committee (IACUC) of Nanjing University (IACUC-2405003). Male SD rats (180-220 g) purchased from Beijing Vital River Laboratory Animal Technology Co., Ltd. (China). Inject 40 of μL complete Freund's adjuvant (Chondrex) into the footpad of the right rear paw for establishing AIA model. Then, these rats were randomly divided into four groups (n = 4). The thickness of right paws was recorded every two days. After immunization for 14 days, the AIA rats were intraarticular injections in ankle joint with PBS, $Fe_{SA+NC}$, BMSCs, and Fe/BMSCs every two days. The rats of control group did not receive any treatment.

**Histomorphometry assessment**

On day 36, the rats were sacrificed and their right ankle tissues were obtained. After fixation in 4% paraformaldehyde, micro-CT scanning was performed for three-dimensional reconstruction of the ankle tissues. The bone histomorphometric parameters were then analyzed using CTAn software.

**Histology assessment**

The ankle tissues were decalcified over four weeks and then stained with hematoxylin and eosin (HE), toluidine bule (TB), and Safranin O/Fast Green (SO).

**In vivo safety assessment**

The main organs (heart, liver, spleen, lung, and kidney) of these rats were collected for HE staining to verify the biocompatibility of nanozymes.

**Statistical analysis**

All data were presented as mean ± standard deviation (SD). One-way ANOVA and Student's t-test were utilized for statistical analyses. Values of $*p < 0.05$, $**p < 0.01$, $***p < 0.001$, and $****p < 0.0001$ were applied to annotate statistical significance.

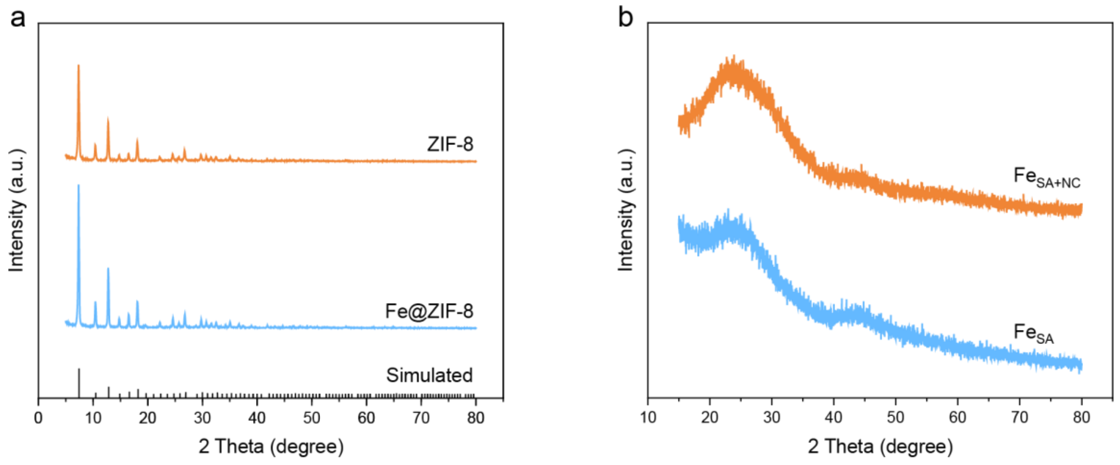


**Figure S1.** XRD patterns of a) ZIF-8 and Fe@ZIF-8, b) $Fe_{SA+NC}$ and $Fe_{SA}$.

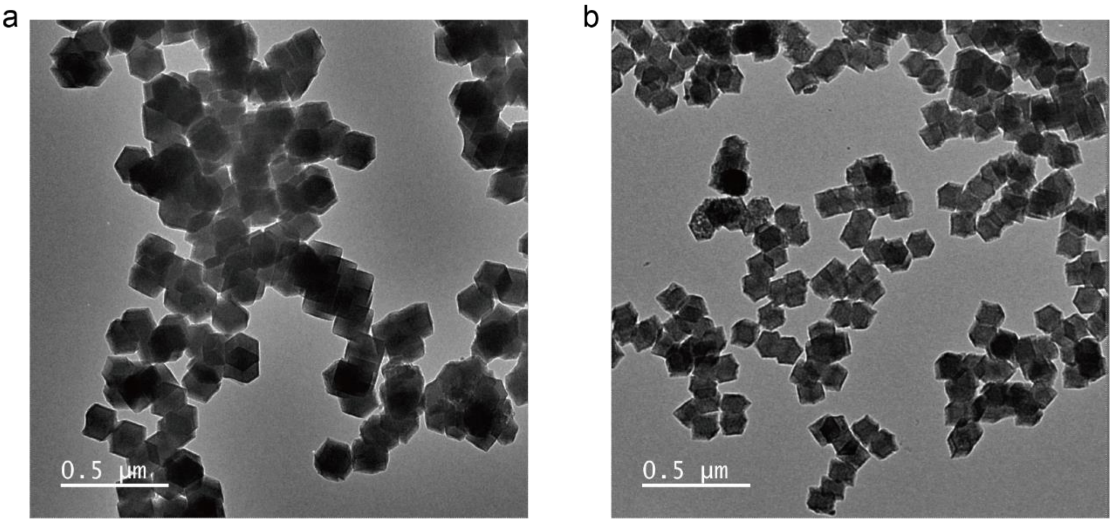


**Figure S2.** TEM images of a) $Fe_{SA+NC}$ and b) $Fe_{SA}$. Scale bars: 0.5 μm.

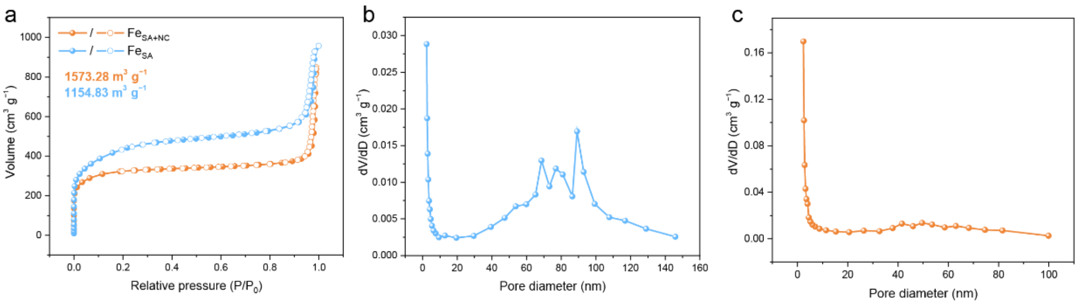


**Figure S3.** a) $N_2$ adsorption/desorption isotherms of $Fe_{SA}$ and $Fe_{SA+NC}$. Pore diameter distribution of b) $Fe_{SA}$ and c) $Fe_{SA+NC}$.

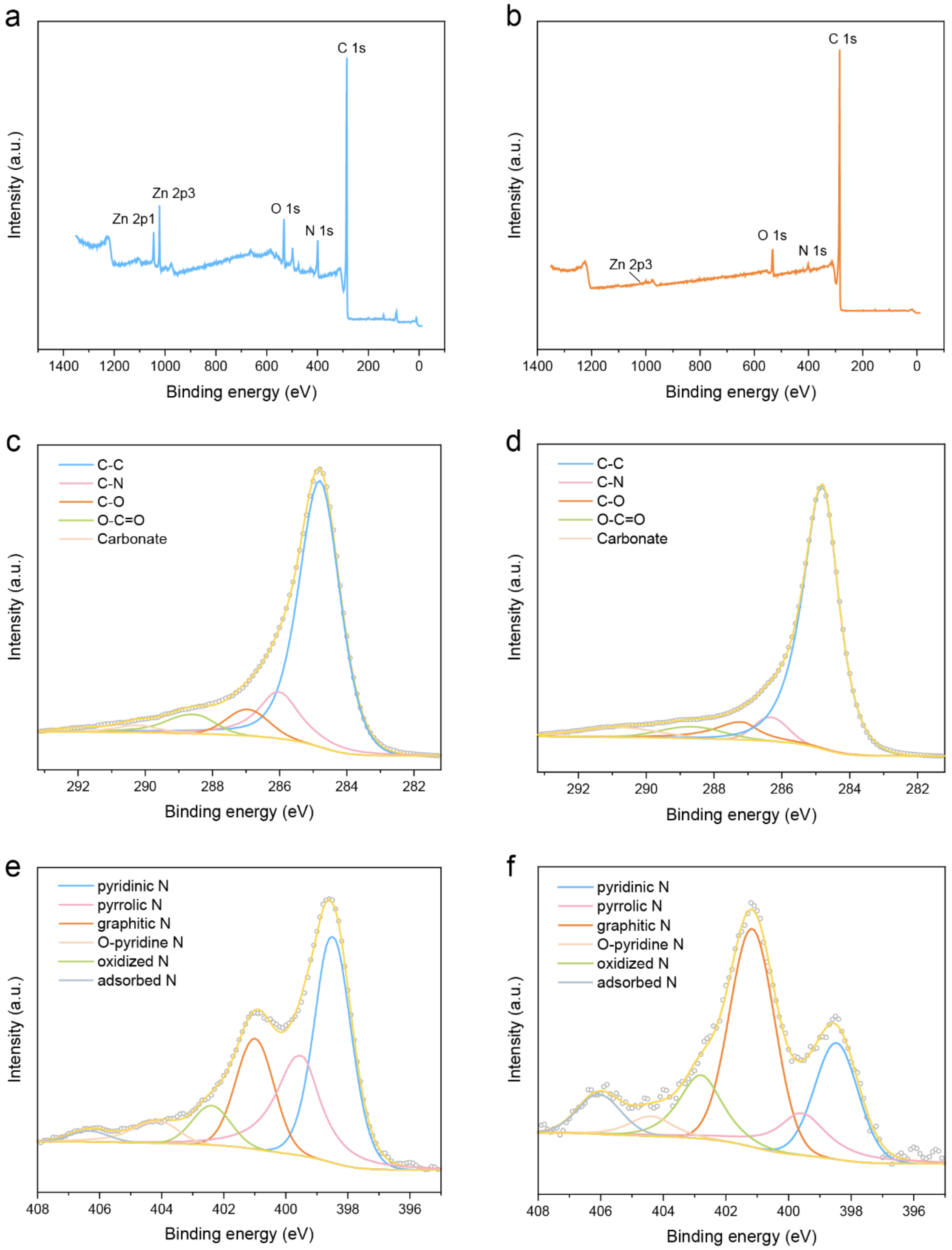


**Figure S4.** a) Survey spectra, high solution c) C 1s and e) N 1s XPS data of $Fe_{SA}$. b) Survey spectra, high solution d) C 1s and f) N 1s XPS data of $Fe_{SA+NC}$.

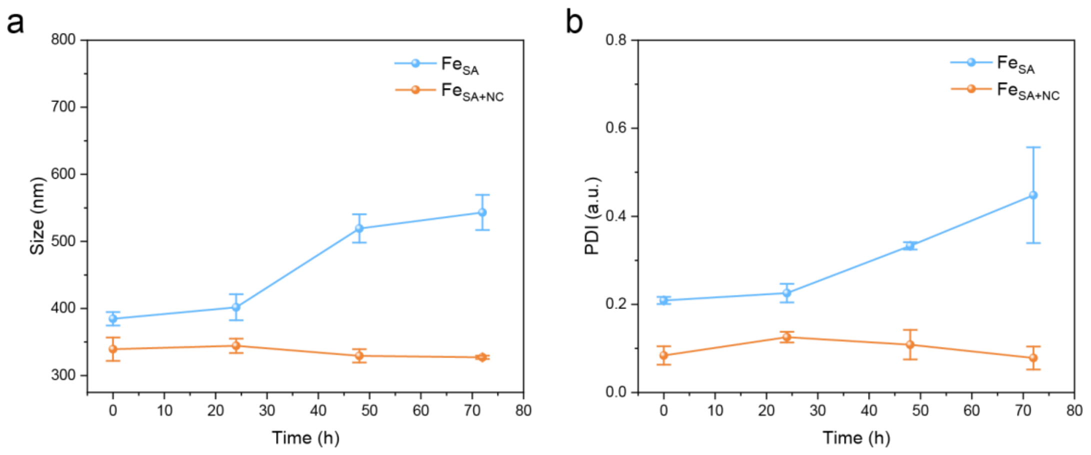


**Figure S5.** Time-dependent changes in a) hydrodynamic diameter and b) polydispersity index (PDI) of $Fe_{SA}$ and $Fe_{SA+NC}$. Data in a) and b) are expressed as mean ± SD, n = 3.

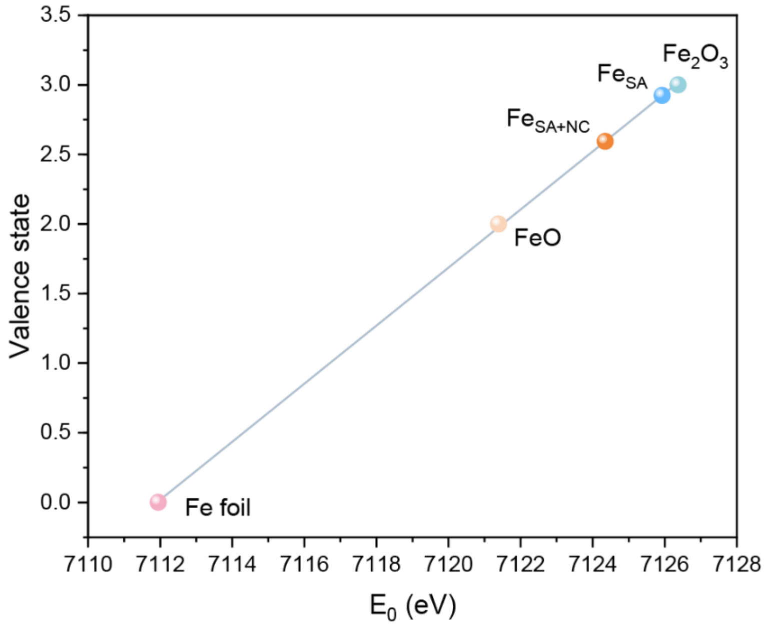


**Figure S6.** Valence state of $Fe_{SA}$ and $Fe_{SA+NC}$.

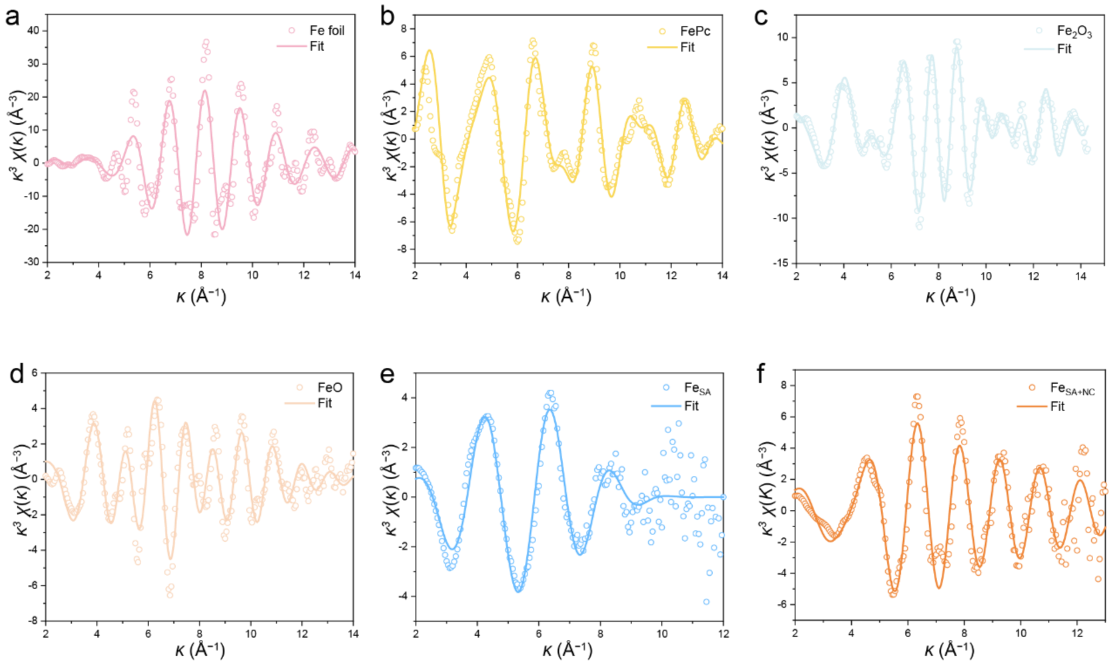


**Figure S7.** FT-EXFAS k space fitting curves of a) Fe foil, b) FePc, c) $Fe_2O_3$, d) FeO, e) $Fe_{SA}$, and f) $Fe_{SA+NC}$.

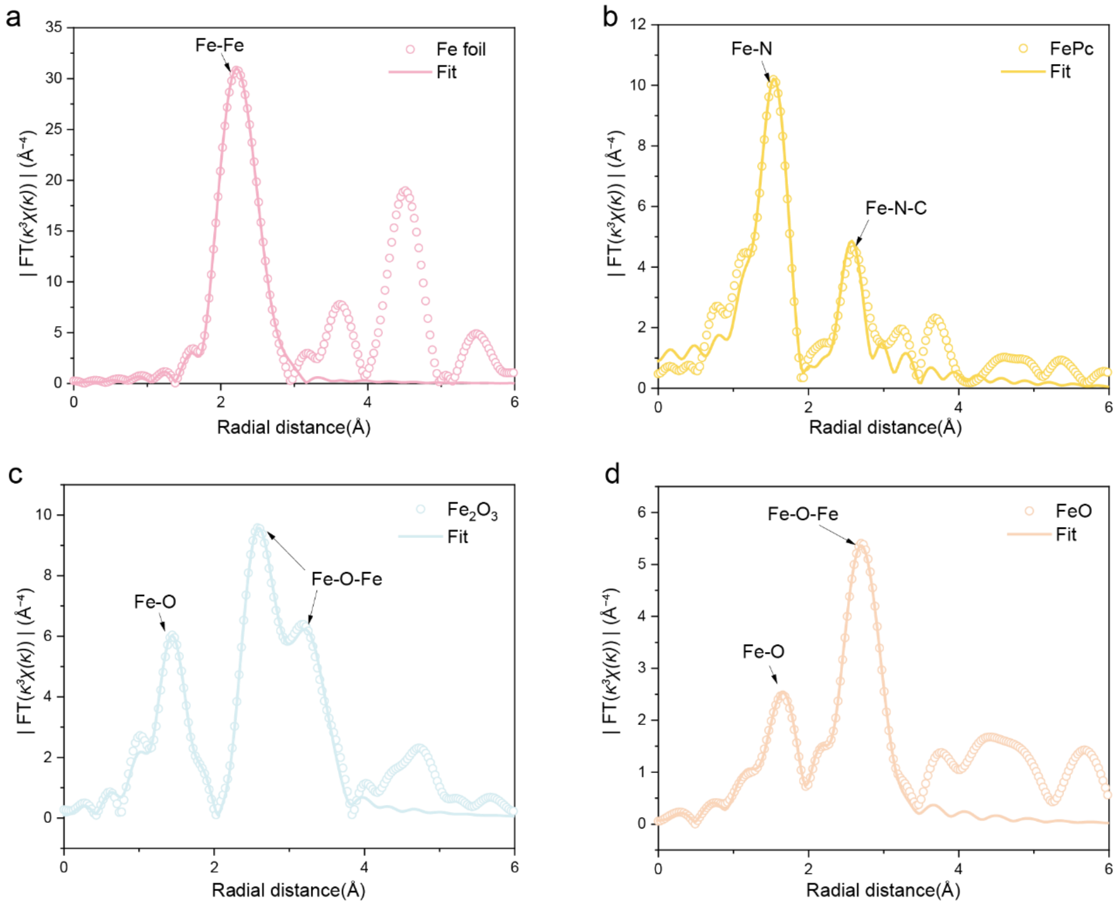


**Figure S8.** FT-EXAFS R space fitting curves a) Fe foil, b) FePc, c) $Fe_2O_3$, and d) FeO.

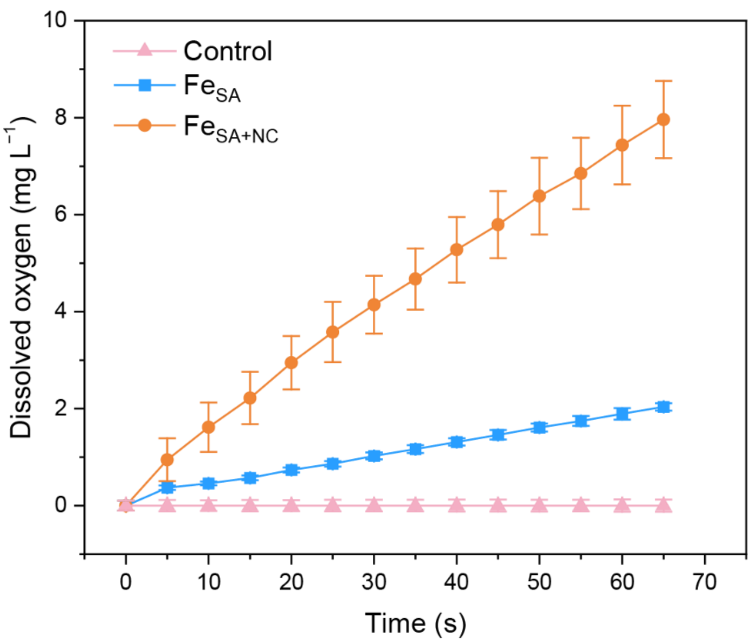


**Figure S9.** Dissolved oxygen produced by $H_2O_2$ solutions containing $Fe_{SA}$ or $Fe_{SA+NC}$. Data are expressed as mean ± SD, n = 3.

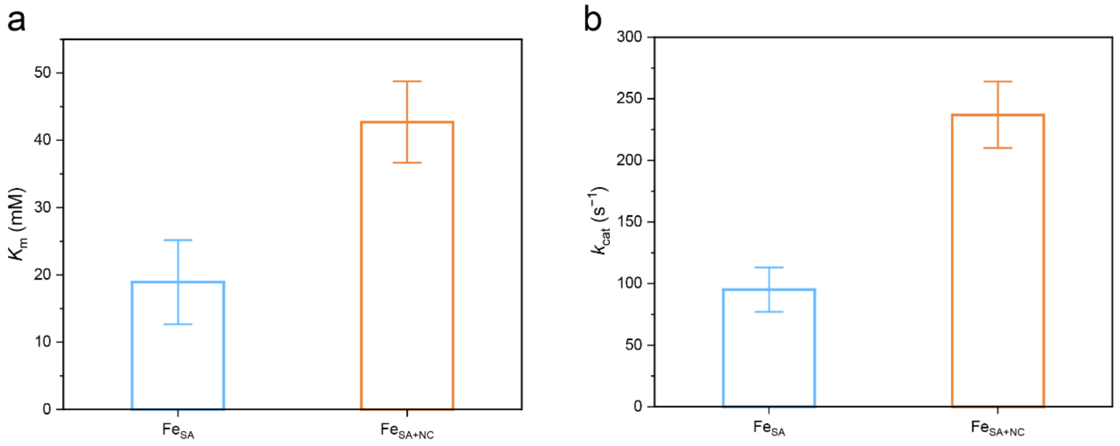


**Figure S10.** a) $K_m$ values and b) $k_{cat}$ values of CAT-like activity of $Fe_{SA}$ and $Fe_{SA+NC}$. Data in a) and b) are expressed as mean ± SD, n = 3.

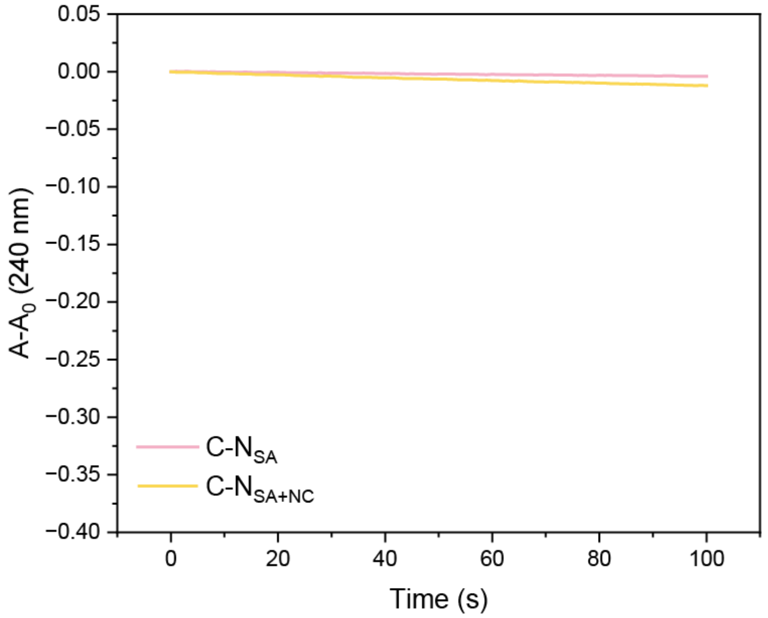


**Figure S11.** CAT-like activities of nitrogen doped carbon carriers.

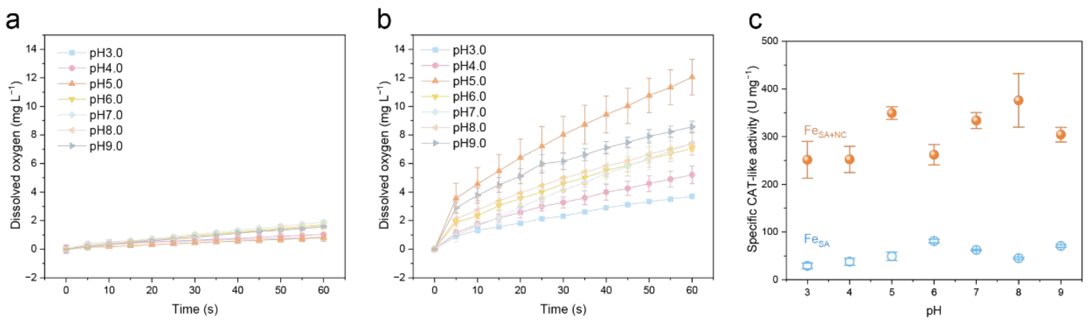


**Figure S12.** pH-dependent CAT-like activity of a) $Fe_{SA}$ and b) $Fe_{SA+NC}$ measured by using a dissolved oxygen meter. c) Specific CAT-like activity of $Fe_{SA}$ and $Fe_{SA+NC}$ under different pH conditions. Data in a–c) are expressed as mean ± SD, n = 3.

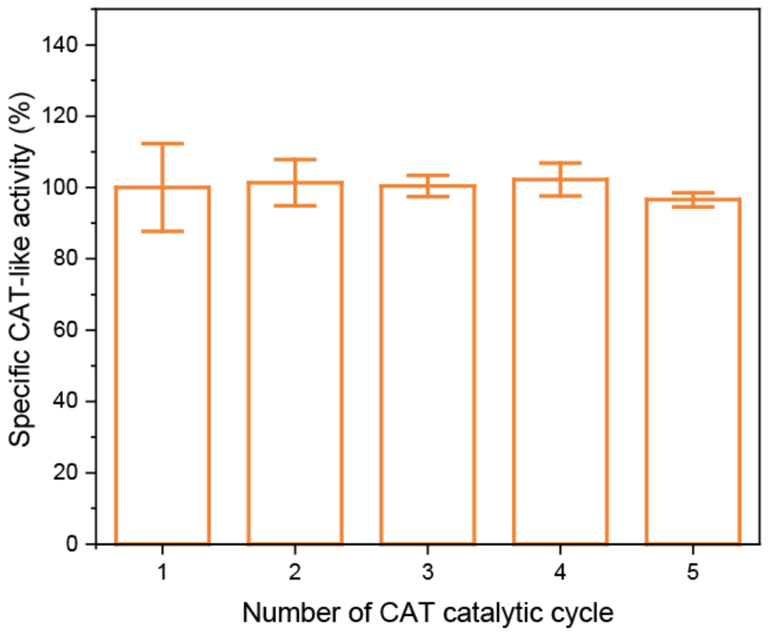


**Figure S13.** Repetitive catalytic performance of $Fe_{SA+NC}$ over five consecutive cycles. Data are expressed as mean ± SD, n = 3.

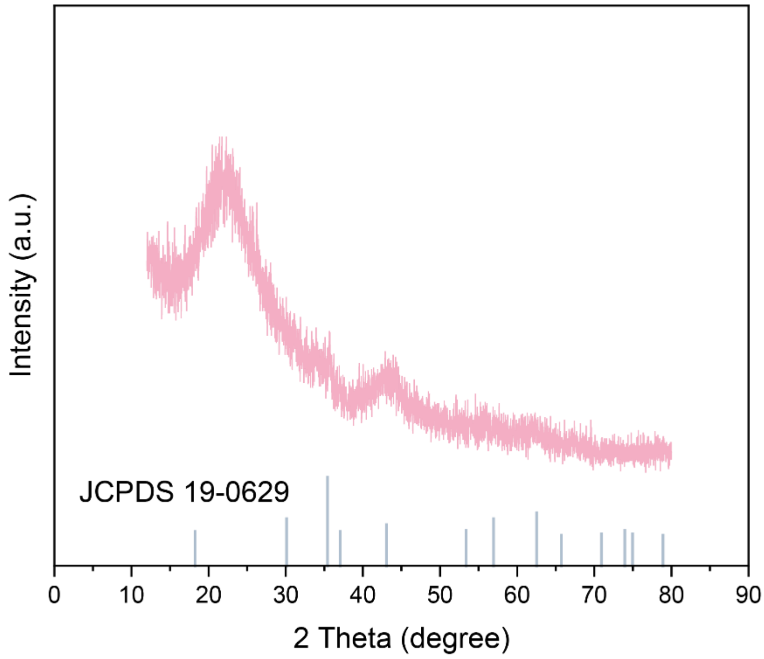


**Figure S14.** XRD pattern of $Fe_{NC}$.

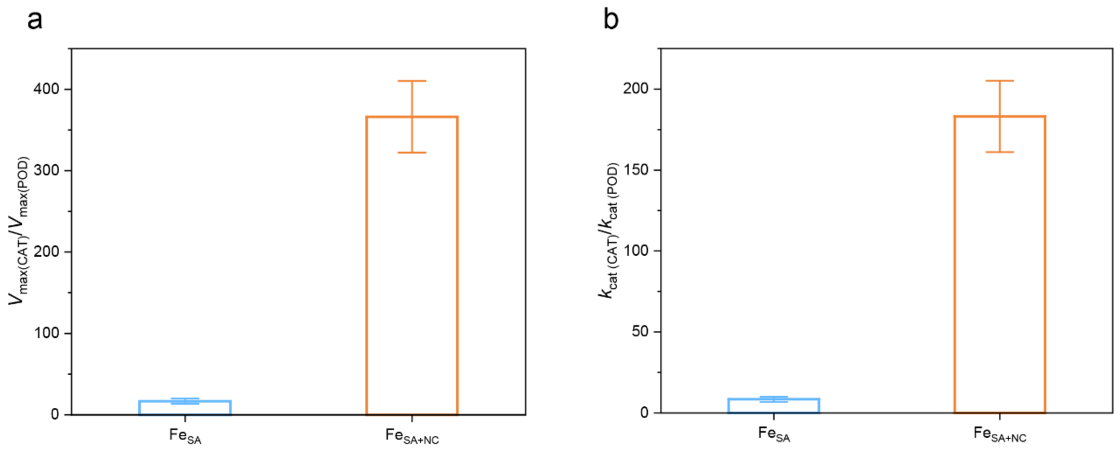


**Figure S15.** Ratios of a) $V_{max}$ and b) $k_{cat}$ for CAT- to POD-like activities of $Fe_{SA}$ and $Fe_{SA+NC}$. Data in a) and b) are expressed as mean ± SD, n = 3.

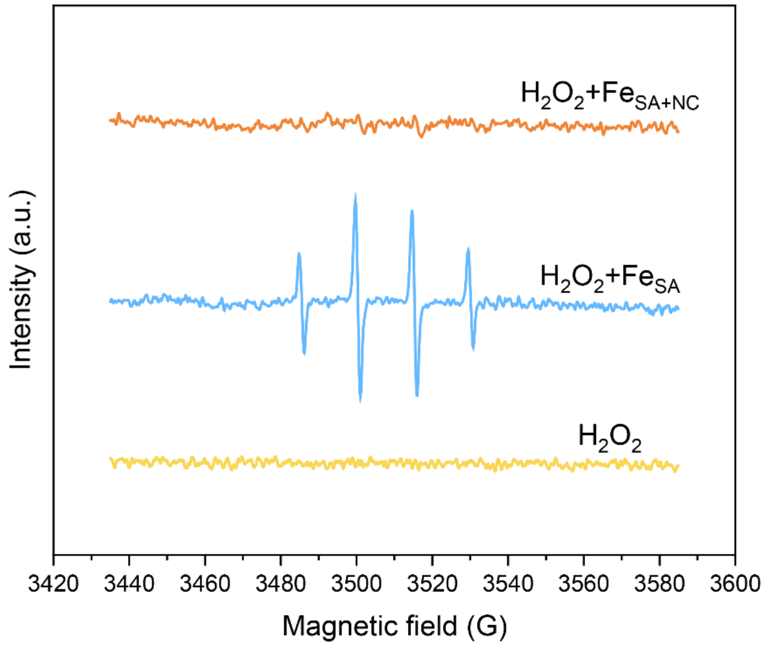


**Figure S16.** EPR spin-trapping of $^{\bullet}$OH generated from $H_2O_2$ after indicated treatments.

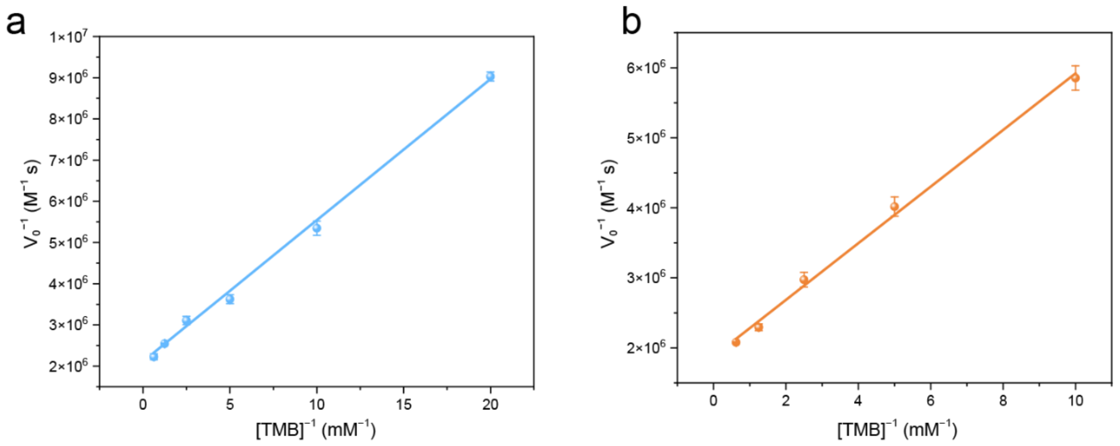


**Figure S17.** Double-reciprocal plots of reciprocal initial velocities versus reciprocal TMB concentrations of a) $Fe_{SA}$ and b) $Fe_{SA+NC}$ for POD-like activity. Data in a) and b) are expressed as mean ± SD, n = 3.

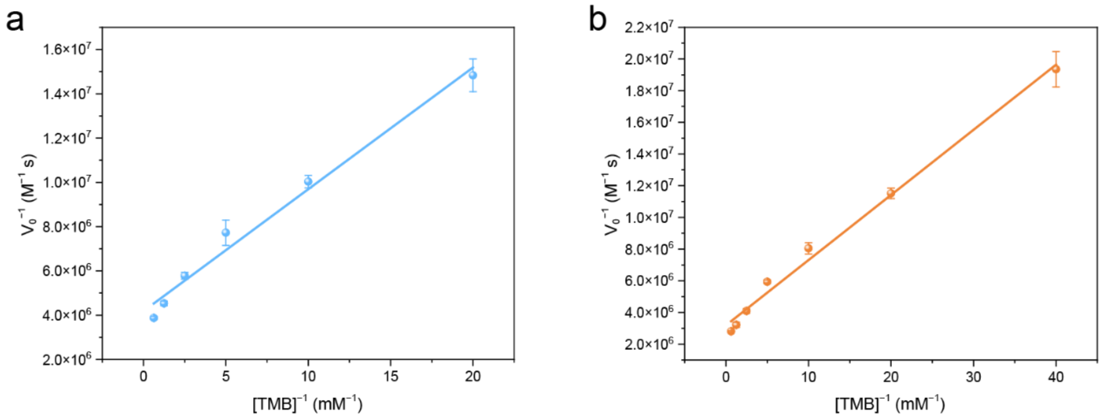


**Figure S18.** Double-reciprocal plots of reciprocal initial velocities versus reciprocal TMB concentrations of a) $Fe_{SA}$ and b) $Fe_{SA+NC}$ for OXD-like activity. Data in a) and b) are expressed as mean ± SD, n = 3.

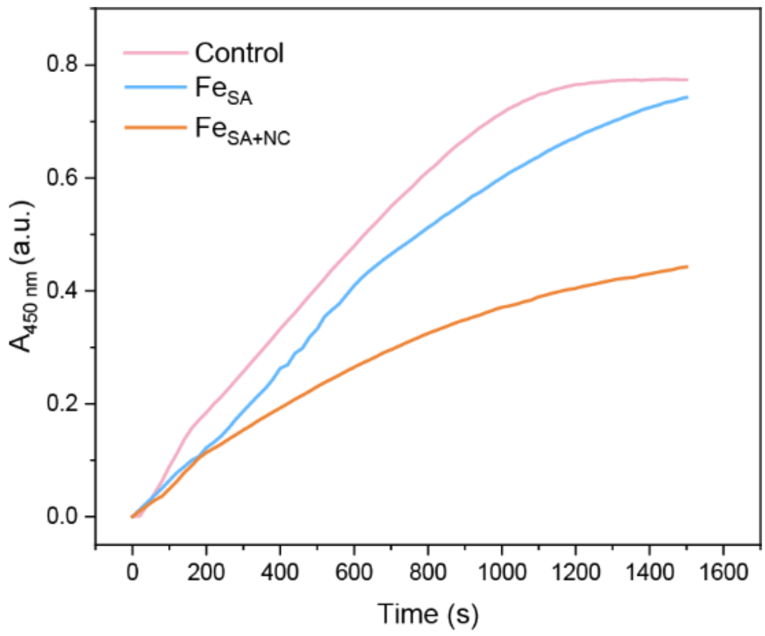


**Figure S19.** Time-dependent absorbance changes at 450 nm of WST-1 during the reaction with $O_2^{\bullet -}$ in the presence of $Fe_{SA}$ or $Fe_{SA+NC}$.

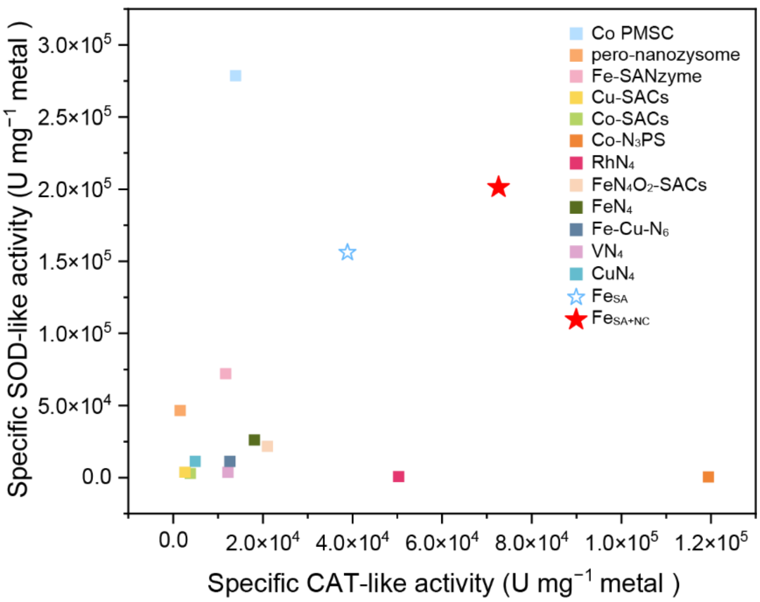


**Figure S20.** Comparison of specific CAT- and SOD-like activities of different SAzymes. Each point represents one catalyst. The red star denotes $Fe_{SA+NC}$ developed in this work. Specific activities are expressed as U $mg^{-1}$ metal atoms. Data and references were listed in Table S8.

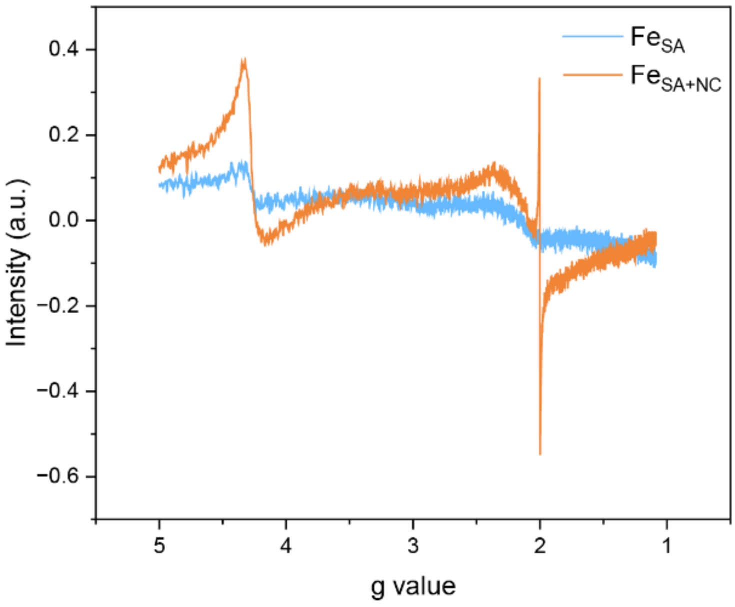


**Figure S21.** EPR spectra of $Fe_{SA}$ and $Fe_{SA+NC}$.

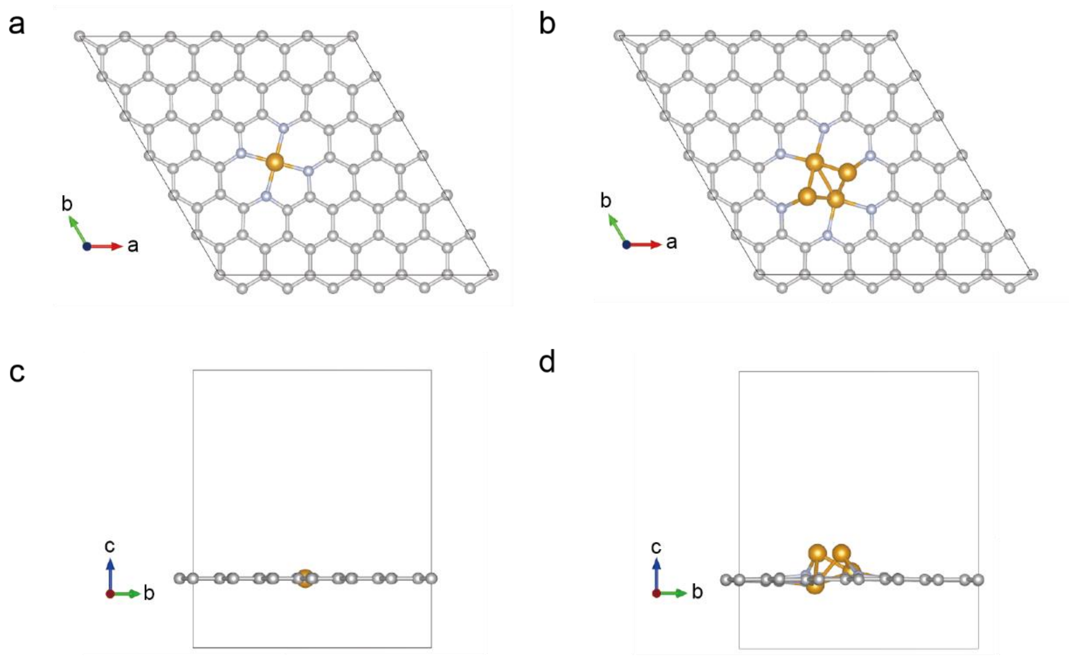


**Figure S22.** The computational model of single-atom Fe ($FeN_4$, a, top view; c, side view) and cluster Fe ($Fe_4N_6$, b, top view; d, side view) catalytic sites. Silver: C atom, light blue: N atom, gold: Fe atom.

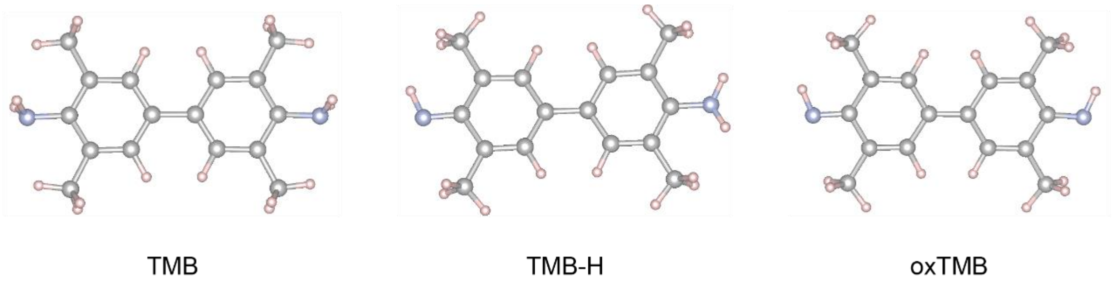


**Figure S23.** The optimized configurations of TMB, TMB-H, and oxTMB, respectively. Silver: C atom, light blue: N atom, light pink: H atom.

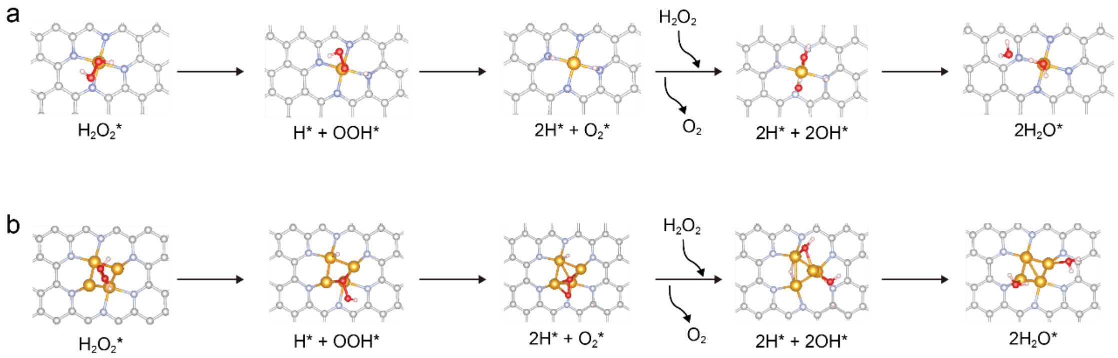


**Figure S24.** The reaction process of CAT-like activity of a) $FeN_4$ and b) $Fe_4N_6$. Silver: C atom, light blue: N atom, gold: Fe atom, light pink: H atom, red: O atom.

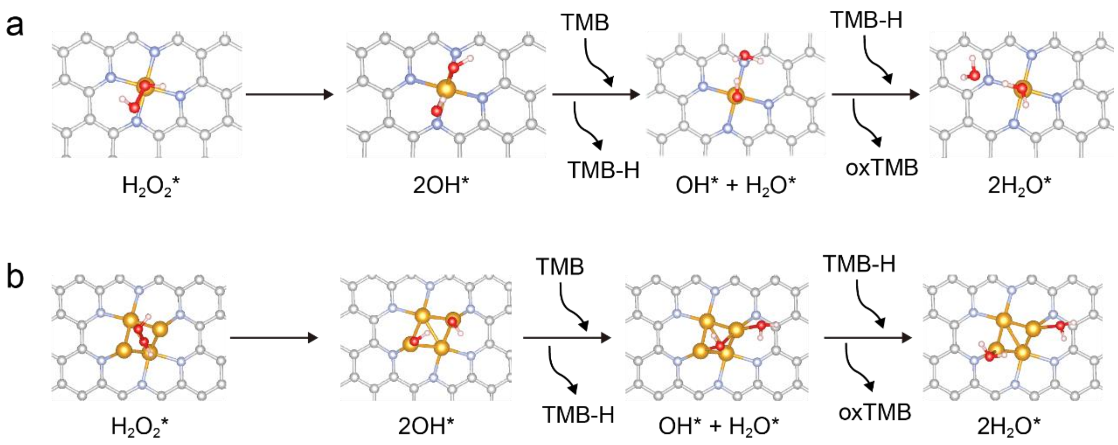


**Figure S25.** The reaction process of POD-like activity of a) $FeN_4$ and b) $Fe_4N_6$. Silver: C atom, light blue: N atom, gold: Fe atom, light pink: H atom, red: O atom.

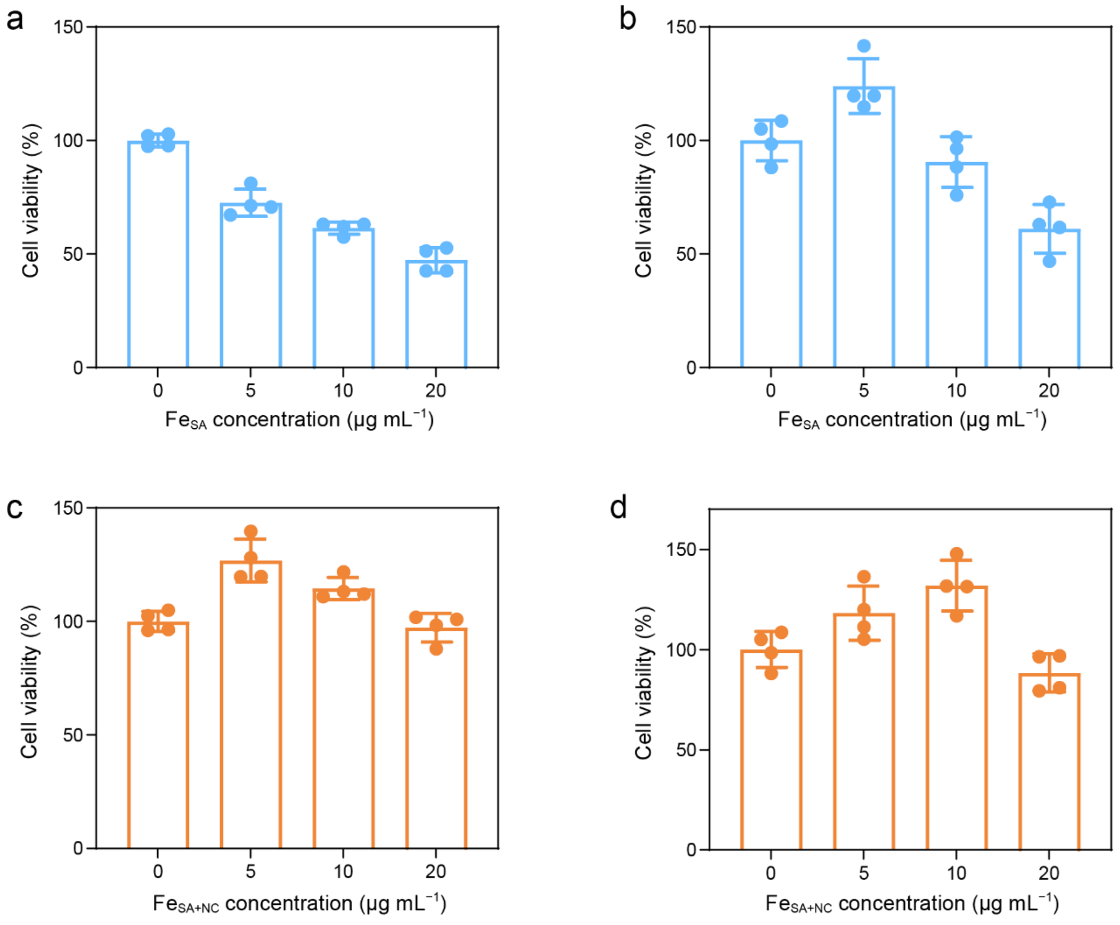


**Figure S26.** Cytotoxicity of Fe$_{SA}$ and Fe$_{SA+NC}$ by using (a, c) RAW264.7 and (b, d) BMSCs. Data in a–d) are expressed as mean ± SD, n = 4.

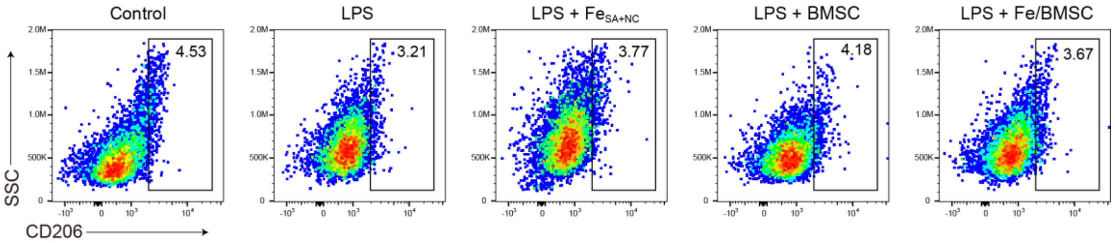


**Figure S27.** Expression of CD206 in macrophages in the co-culture system evaluated by flow cytometry.

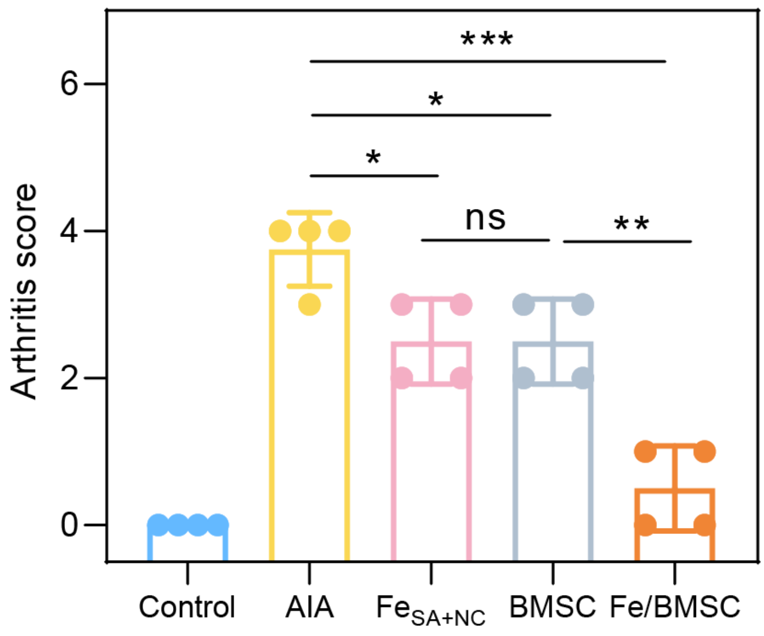


**Figure S28.** Average arthritis scores for different groups. Data are expressed as mean ± SD, n = 4.

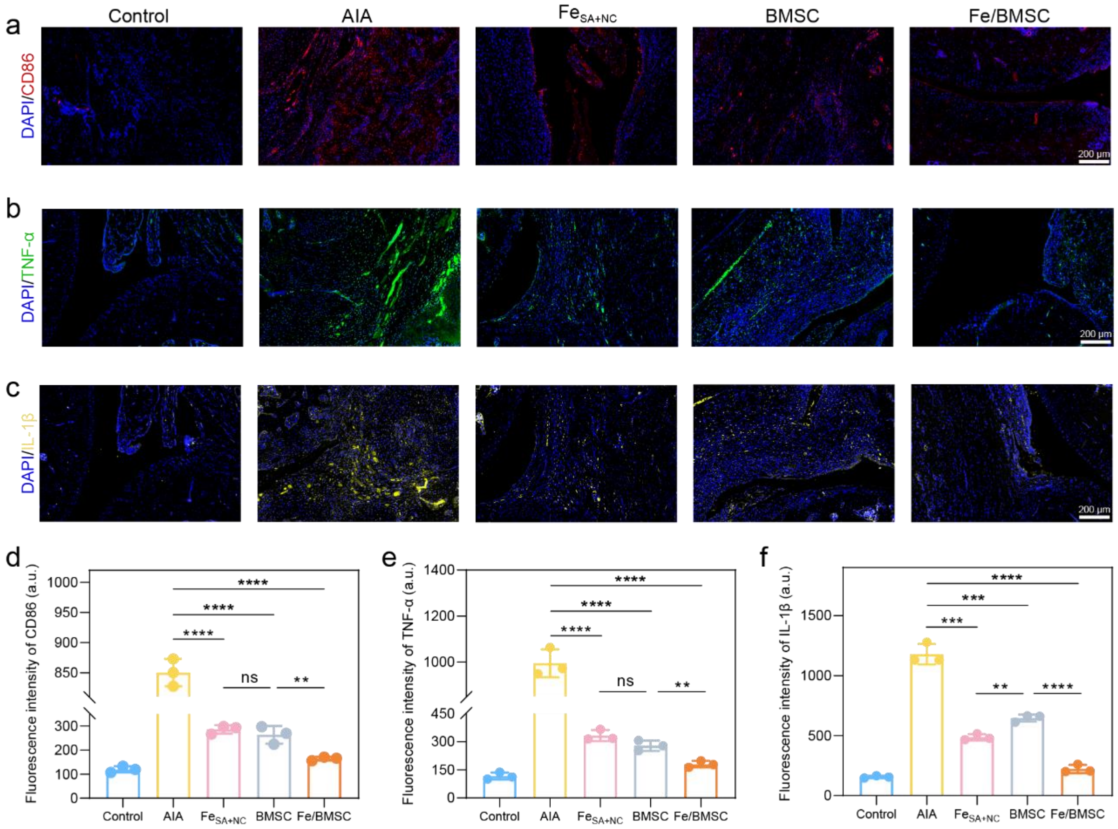


**Figure S29.** a–c) Immunofluorescence staining and d–f) quantitative analysis of CD86, TNF-α, and IL-1β in the normal and inflamed joints after different treatments. Data in d–f) are expressed as mean ± SD, n = 3. Scale bars: 200 μm.

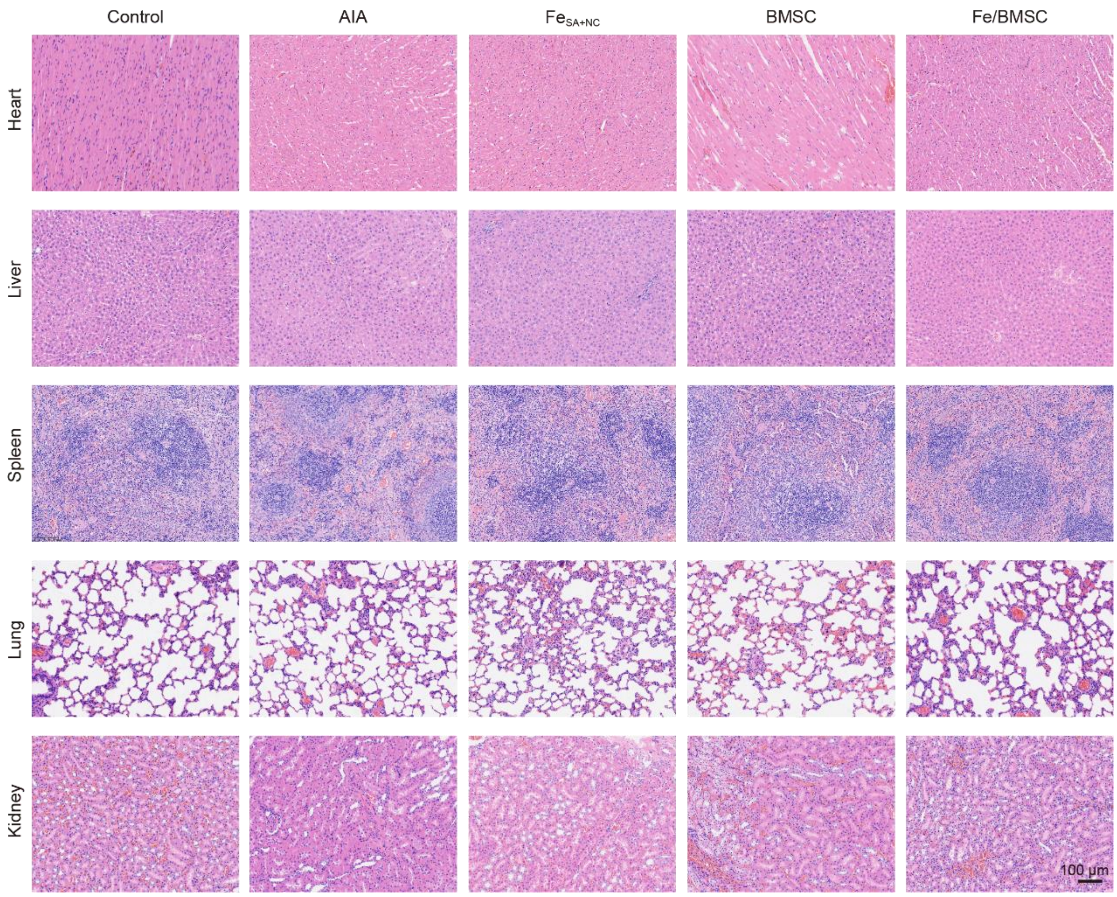


**Figure S30.** Histological morphologies of major organs of healthy control and AIA model rats after different treatments at the end of therapy. Scale bar: 100 μm.

**Table S1.** BET surface areas and pore volumes of Fe–N–C nanozymes.

| Sample | Surface area $m^2\ g^{-1}$ | Pore volume $cm^3\ g^{-1}$ |
|---|---|---|
| $Fe_{SA}$ | 1154.83 | 0.85 |
| $Fe_{SA+NC}$ | 1573.28 | 0.91 |

**Table S2.** Fitting results of C 1s XPS spectra for Fe–N–C nanozymes (at. %).

| Sample | C=C | C–N/C–C | C–O | O–C=O | Carbonate |
|---|---|---|---|---|---|
| $Fe_{SA}$ | 70.46 | 14.90 | 6.50 | 6.14 | 1.99 |
| $Fe_{SA+NC}$ | 76.35 | 5.69 | 7.24 | 5.25 | 5.46 |

**Table S3.** Fitting results of N 1s XPS spectra for Fe–N–C nanozymes (at. %).

| Sample | Pyridinic N | Pyrrolic N | Graphitic N | O-pyridine N | Oxidized N | Adsorbed N |
|---|---|---|---|---|---|---|
| $Fe_{SA}$ | 40.28 | 25.71 | 19.79 | 7.05 | 5.21 | 1.96 |
| $Fe_{SA+NC}$ | 21.68 | 12.52 | 41.60 | 3.94 | 12.86 | 7.41 |

**Table S4.** Fe content in $Fe_{SA}$ and $Fe_{SA+NC}$ measured by ICP (wt %).

| Sample | Content 1 | Content 2 | Content 3 | Average |
|---|---|---|---|---|
| $Fe_{SA}$ | 0.13 | 0.17 | 0.18 | 0.16 |
| $Fe_{SA+NC}$ | 0.44 | 0.49 | 0.44 | 0.46 |

**Table S5.** EXAFS data fitting results of Fe–N–C nanozymes.

| Sample | Path | $CN^a$ | $R$(Å)$^b$ | $\sigma^2$ (Å$^2$)$^c$ | $\Delta E_0$(eV)$^d$ | $R$ factor |
|---|---|---|---|---|---|---|
| Fe K-edge ($S_0^2$ = 0.765) | | | | | | |
| Fe foil | Fe–Fe | 8.0* | 2.474 ± 0.006 | 0.0050 | 7.5 ± 1.1 | 0.0016 |
| | Fe–Fe | 6.0* | 2.856 ± 0.006 | 0.0069 | | |
| FeO | Fe–O | 6.1 ± 0.7 | 2.141 ± 0.011 | 0.0138 | 4.9 ± 1.1 | 0.0010 |
| | Fe–O–Fe | 12.8 ± 1.0 | 3.075 ± 0.003 | 0.0090 | 2.7 ± 0.5 | |
| FePc | Fe–N | 4.0 ± 0.2 | 1.940 ± 0.006 | 0.0051 | 9.8 ± 1.4 | 0.0184 |
| | Fe–N–C | 9.4 ± 1.1 | 2.962 ± 0.010 | 0.0033 | | |
| $Fe_2O_3$ | Fe–O | 3.8 ± 0.2 | 1.942 ± 0.010 | 0.0042 | −1.8 ± 1.7 | 0.0034 |
| | Fe–O | 2.4 ± 0.3 | 2.111 ± 0.017 | | | |
| | Fe–O–Fe | 4.7 ± 0.2 | 2.957 ± 0.006 | 0.0060 | −0.9 ± 1.1 | |
| | Fe–O–Fe | 3.7 ± 0.3 | 3.385 ± 0.009 | | | |
| | Fe–O–Fe | 4.8 ± 0.3 | 3.691 ± 0.007 | | | |
| $Fe_{SA}$ | Fe–N | 4.1 ± 0.3 | 1.989 ± 0.006 | 0.0072 | 1.1 ± 3.0 | 0.0105 |
| $Fe_{SA+NC}$ | Fe–N | 3.9 ± 0.2 | 1.994 ± 0.016 | 0.0147 | 2.4 ± 2.0 | 0.0021 |
| | Fe–Fe | 2.7 ± 0.1 | 2.510 ± 0.004 | 0.0057 | −4.1 ± 0.9 | |

$^a$*CN*, coordination number; $^b$*R*, the distance between absorber and backscatter atoms; $^c\sigma^2$, the Debye Waller factor value; $^d\Delta E_0$, inner potential correction to account for the difference in the inner potential between the sample and the reference compound; *R* factor indicates the goodness of the fit. $S0^2$ was fixed to 0.765, according to the experimental EXAFS fit of Fe foil by fixing *CN* as the known crystallographic value. * This value was fixed during EXAFS fitting, based on the known structure of Fe. Fitting conditions: *k* range: 3.0–11.0; *R* range: 1.0–3.0; fitting space: R space; *k*-weight = 3. A reasonable range of EXAFS fitting parameters: $0.800 < S_0^2 < 1.000$; $CN > 0$; $\sigma^2 > 0$ Å$^2$; $|\Delta E_0| < 15$ eV; *R* factor < 0.02.

**Table S6.** Comparison of the kinetic parameters with $H_2O_2$ as the substrate.

| Substrate | | $Fe_{SA}$ | | $Fe_{SA+NC}$ | |
|---|---|---|---|---|---|
| | | catalase | peroxidase | catalase | peroxidase |
| $H_2O_2$ | $K_m$ (mM) | 18.92 | 12.08 | 42.70 | 1.32 |
| | $V_{max}$ (mM s$^{-1}$) | $5.45 \times 10^{-2}$ | $3.23 \times 10^{-3}$ | 0.39 | $1.07 \times 10^{-3}$ |
| | $k_{cat}$ (s$^{-1}$) | 95.12 | 11.27 | 237.00 | 1.29 |
| | Specific activity (U mg$^{-1}$) | 62.39 | 60.39 | 333.79 | 38.49 |

**Table S7.** Comparison of the kinetic parameters with TMB as the substrate.

| Substrate | | $Fe_{SA}$ | | $Fe_{SA+NC}$ | |
|---|---|---|---|---|---|
| | | peroxidase | oxidase | peroxidase | oxidase |
| TMB | $K_m$ (mM) | 0.16 | 0.18 | 0.23 | 0.21 |
| | $V_{max}$ (mM s$^{-1}$) | $4.73 \times 10^{-4}$ | $2.7 \times 10^{-4}$ | $5.5 \times 10^{-4}$ | $3.8 \times 10^{-4}$ |
| | $k_{cat}$ (s$^{-1}$) | 1.65 | 0.95 | 0.67 | 0.46 |
| | Specific activity (U mg$^{-1}$) | 60.39 | 1.30 | 38.49 | 1.78 |

**Table S8.** Comparison of specific CAT- and SOD-like activities (U mg$^{-1}$) of reported M–N–C type nanozymes normalized to metal content.

| Sample | Content (wt%) | CAT (U mg$^{-1}$) | SOD (U mg$^{-1}$) | Reference |
|---|---|---|---|---|
| Fe in $Fe_1$/NC-900 | 0.28 | $1.03 \times 10^4$ | / | [4] |
| Fe in pero-nanozysome | 2.7 | $0.15 \times 10^4$ | $4.66 \times 10^4$ | [5] |
| Fe in Fe-SANzyme | 0.45 | $1.17 \times 10^4$ | $7.21 \times 10^4$ | [6] |
| Ir in Ir NC SAzymes | 0.59 | $0.97 \times 10^4$ | / | [7] |
| Fe in $Fe_2$-SAzyme | 0.41 | $1.68 \times 10^4$ | / | [8] |
| Co in Co PMSC | 2.02 | $1.39 \times 10^4$ | $27.87 \times 10^4$ | [9] |
| Co in Co-$N_3$PS | / | $11.94 \times 10^4$ | / | [10] |
| Rh in $RhN_4$ | 0.19 | $5.03 \times 10^4$ | $0.06 \times 10^4$ | [11] |
| V in $VN_4$ | 0.22 | $1.22 \times 10^4$ | $0.08 \times 10^4$ | [11] |
| Fe in $FeN_4$ | 1.40 | $1.81 \times 10^4$ | $0.37 \times 10^4$ | [11] |
| Cu in $CuN_4$ | 1.39 | $0.49 \times 10^4$ | $2.61 \times 10^4$ | [11] |
| Fe-Cu in Fe–Cu–$N_6$ | 1.10 | $1.26 \times 10^4$ | $1.13 \times 10^4$ | [11] |
| Co in Co-SACs | / | $0.38 \times 10^4$ | $0.29 \times 10^4$ | [12] |
| Cu in Cu-SACs | / | $0.26 \times 10^4$ | $0.38 \times 10^4$ | [12] |
| Fe in $FeN_4O_2$-SACs | 0.43 | $2.1 \times 10^4$ | $2.18 \times 10^4$ | [12] |
| Fe in $Fe_{SA}$ | 0.16 | $3.89 \times 10^4$ | $15.61 \times 10^4$ | This work |

| | | | | |
|---|---|---|---|---|
| Fe in $Fe_{SA+NC}$ | 0.46 | $7.26 \times 10^4$ | $20.13 \times 10^4$ | This work |

### References


[1] Kresse, G.; Joubert, D. From Ultrasoft Pseudopotentials to the Projector Augmented-Wave Method. *Physical Review B* **1999**, *59*, 1758-1775.

[2] Perdew, J. P.; Burke, K.; Ernzerhof, M. Generalized Gradient Approximation Made Simple. *Physical Review Letters* **1996**, *77*, 3865-3868.

[3] Grimme, S.; Antony, J.; Ehrlich, S.; Krieg, H. A Consistent and Accurate *Ab Initio* Parametrization of Density Functional Dispersion Correction (DFT-D) for the 94 Elements H-Pu. *The Journal of Chemical Physics* **2010**, *132*, 154104.

[4] Gao, X.; Wei, H.; Ma, W.; Wu, W.; Ji, W.; Mao, J.; Yu, P.; Mao, L. Inflammation-Free Electrochemical in Vivo Sensing of Dopamine with Atomic-Level Engineered Antioxidative Single-Atom Catalyst. *Nature Communications* **2024**, *15*, 7915.

[5] Xi, J.; Zhang, R.; Wang, L.; Xu, W.; Liang, Q.; Li, J.; Jiang, J.; Yang, Y.; Yan, X.; Fan, K.; Gao, L. A Nanozyme-Based Artificial Peroxisome Ameliorates Hyperuricemia and Ischemic Stroke. *Advanced Functional Materials* **2021**, *31*, 2007130.

[6] Zhang, R.; Xue, B.; Tao, Y.; Zhao, H.; Zhang, Z.; Wang, X.; Zhou, X.; Jiang, B.; Yang, Z.; Yan, X.; Fan, K. Edge-Site Engineering of Defective Fe-$N_4$ Nanozymes with Boosted Catalase-Like Performance for Retinal Vasculopathies. *Advanced Materials* **2022**, *34*, 2205324.

[7] Wang, Z.; Wang, W.; Wang, J.; Wang, D.; Liu, M.; Wu, Q.; Hu, H. Single-Atom Catalysts with Ultrahigh Catalase-Like Activity Through Electron Filling and Orbital Energy Regulation. *Advanced Functional Materials* **2023**, *33*, 2209560.

[8] Zhang, H.; Wang, P.; Zhang, J.; Sun, Q.; He, Q.; He, X.; Chen, H.; Ji, H. Boosting the Catalase-Like Activity of SAzymes via Facile Tuning of the Distances between Neighboring Atoms in Single-Iron Sites. *Angewandte Chemie International Edition* **2024**, *63*, e202316779.

[9] Cao, F.; Zhang, L.; You, Y.; Zheng, L.; Ren, J.; Qu, X. An Enzyme-Mimicking Single-Atom Catalyst as an Efficient Multiple Reactive Oxygen and Nitrogen Species Scavenger for Sepsis Management. *Angewandte Chemie International Edition* **2020**, *59*, 5108-5115.

[10] Chen, Y.; Jiang, B.; Hao, H.; Li, H.; Qiu, C.; Liang, X.; Qu, Q.; Zhang, Z.; Gao, R.; Duan, D.; Ji, S.; Wang, D.; Liang, M. Atomic-Level Regulation of Cobalt Single-Atom Nanozymes: Engineering High-Efficiency Catalase Mimics. *Angewandte Chemie International Edition* **2023**, *62*, e202301879.

[11] Zhang, S.; Li, Y.; Sun, S.; Liu, L.; Mu, X.; Liu, S.; Jiao, M.; Chen, X.; Chen, K.; Ma, H.; Li, T.; Liu, X.; Wang, H.; Zhang, J.; Yang, J.; Zhang, X. Single-Atom Nanozymes Catalytically Surpassing Naturally Occurring Enzymes as Sustained Stitching for Brain Trauma. *Nature Communications* **2022**, *13*, 4744.

[12] Lu, X.; Kuai, L.; Huang, F.; Jiang, J.; Song, J.; Liu, Y.; Chen, S.; Mao, L.; Peng, W.; Luo, Y.; Li, Y.; Dong, H.; Li, B.; Shi, J. Single-Atom Catalysts-Based Catalytic ROS Clearance for Efficient Psoriasis Treatment and Relapse Prevention via Restoring ESR1. *Nature Communications* **2023**, *14*, 6767.
